# Causal evidence of network communication in whole-brain dynamics through a multiplexed neural code

**DOI:** 10.1101/2020.06.09.142695

**Authors:** Piergiorgio Salvan, Alberto Lazari, Diego Vidaurre, Francesca Mandino, Heidi Johansen-Berg, Joanes Grandjean

## Abstract

An important question in neuroscience is how local activity can be flexibly and selectively routed across the brain network. A proposed mechanism to flexibly route information is frequency division multiplexing: selective readout can be achieved by segregating the signal into non-overlapping frequency bands. Here, in wild-type mice and in a transgenic model (3xTgAD) of Alzheimer’s Disease (AD), we use optogenetic activation of the entorhinal cortex, concurrent whole-brain fMRI, and hidden Markov modeling. We demonstrate how inducing neuronal spiking with different theta frequencies causes spatially distinct states of brain network dynamics to emerge and to preferentially respond to one frequency, showing how selective information streams can arise from a single neuronal source of activity. This theta modulation mechanism, however, is impaired in the AD model. This work demonstrates that neuronal multiplexing is a sufficient mechanism to enable flexible brain network communication, and provides insight into the aberrant mechanisms underlying cognitive decline.

## Introduction

The transient synchronization of neuronal spiking, a state in which the activity of a neuronal population is periodically synchronized either within or across brain regions, is an ubiquitous property in brain networks. Most importantly, the spatiotemporal features of these network oscillations are known to vary with cognitive tasks, suggesting a central role in neural coding ^1–3^. A prominent hypothesis on the computational role of oscillatory brain network dynamics is that they function to control information flow through long-range anatomical pathways ^4^. Because distinct behavioral tasks require different combinations of highly specialised regions to work together, routing information via network oscillations would have a crucial role in supporting flexible and selective brain network communication and thus efficient behavior. Recent work has shown evidence of phase-coupling frequency-specific signatures of organization in spontaneous human brain activity *at rest* measured via MEG, possibly reflecting the functional specialisation of distinct circuits at different intrinsic timescales ^5^. These spontaneous spatiotemporal patterns of network oscillations unfold dynamically across time, and may be referred to as the “communication dynamics” of the system ^6^.

However, the mechanisms that would allow *flexible* communication dynamics to emerge in a networked system remain unknown. Although network dynamics may originate from multiple sources of oscillatory activity, it is still unclear whether single-area neuronal spiking can also give rise to multiple states of network dynamics. And furthermore, it remains unclear what role brain topology plays in shaping network dynamics.

Here, we analyzed blood oxygen level dependent (BOLD) fMRI fluctuations of the mouse whole brain, both at rest and during optogenetic stimulation of the EC. We used hidden Markov models (HMM) that, in a completely data-driven fashion, characterize a discrete set of HMM states: transient, repeating patterns of BOLD brain signal amplitude and functional connectivity (**Fig. 1**). In other words, each state represents a unique brain network state of distinct activity and functional connectivity that recurs at different points in time during the scan ^7^. Using HMM, we find that spontaneous BOLD fMRI brain activity in mice is organized in a hierarchical fashion that is similar to humans. We then combine fMRI with optogenetic activation of the EC to demonstrate that localized neuronal spiking is sufficient to cause distinct network states, with different short-scale and long-scale temporal trajectories. Crucially, these network states exhibit preferential responses to different frequencies in neuronal spiking, and this frequency-specific response is impaired in a transgenic model of Alzheimer’s disease. Together, our results demonstrate the biological validity and reproducibility of our fMRI-HMM approach in mice and highlight the relevance of this approach to study the cell-type specific mechanisms underlying neuronal coding in brain network communication.

**Figure 1.**
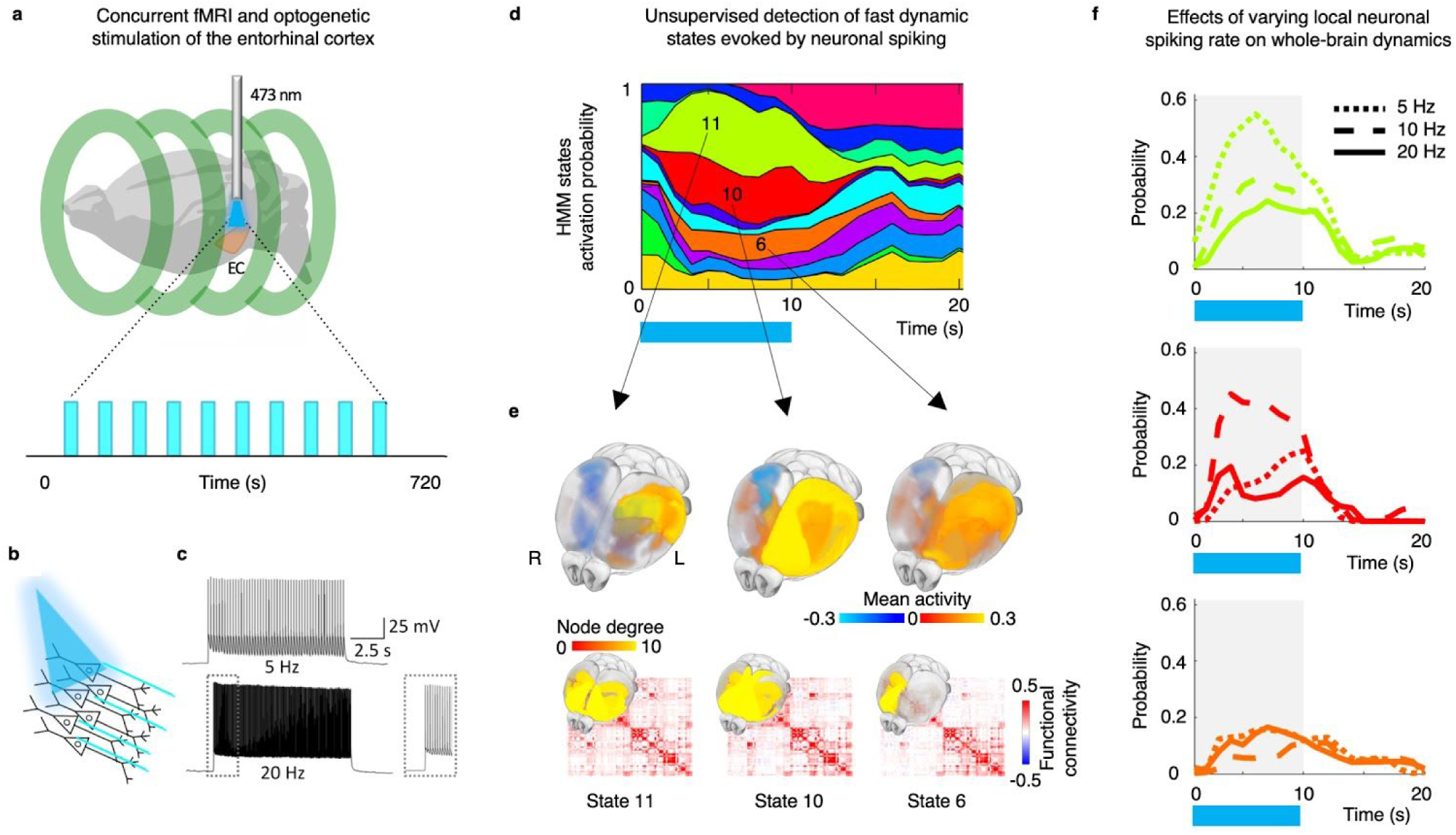
Hidden Markov modeling brain network dynamics during optogenetic stimulation of excitatory neurons in the EC. **a)** Lightly-anesthetized mice under mechanical ventilation underwent high-field functional MRI acquisition runs, concurrently with 10 optogenetics photostimulation blocks of 10 s interleaved with 50 s inter-stimuli intervals per run. Photostimulation was conducted in the left lateral entorhinal cortex (EC) with frequency range 5 to 20 Hz pseudo-randomized between runs. **b)** Photostimulation of EC neurons elicited a faithful response to both 5 and 20 Hz stimulation pulses, here shown in wild-type mice, as confirmed in ex-vivo slice preparation **(c). d)** Whole-brain fMRI BOLD fluctuations in response to optogenetic stimulation were modelled through Hidden Markov models (HMMs), characterizing a discrete number of dynamic brain networks, or states. Temporally, they are represented by a specific state time course that, for each subject, indicates when each state is active (activation probability). Showing WT-ChR2 group average HMM states activation probability time-locked to the optogenetic stimulation (shown by blue bar underneath). **e**) Spatially, each state is characterized by BOLD mean activation (amplitude) and functional connectivity matrix (also represented as network degree). **f**) The effect of varying EC neuronal spiking rate on brain network dynamics was then assessed.

## Results

Oscillations result from the dynamic interaction between cellular and circuit properties. At microscopic scales, oscillatory synchronizations are constrained by axon conduction and synaptic delays; at macroscopic scales, oscillatory synchronizations are thought to be influenced by the architectural properties of the brain network (for review: ^8^). This complex interplay between oscillatory patterns and brain network topology may, therefore, allow neuronal signalling to *selectively* propagate along distinct subdivisions of anatomical projections or circuits ^6^, giving rise to multiple temporal and spatial scales ^8^ or information streams ^4^. From a modeling perspective, it is possible to utilize the sequential nature of BOLD fMRI data in order to characterize transient network states in brain hemodynamic activity. HMMs are well-suited because they provide a time-point-by-time-point description of network activity at the single-trial level, facilitating the characterisation of how such responses may evolve across time ^7^ or in response to perturbations such as stimulation.

While HMMs provide a novel way of interrogating information within fMRI data, their use has not been previously applied to species other than humans. Here, using HMMs, we show that spontaneous hemodynamic fluctuations in the mouse brain at rest are temporally organized in a series of states that mirror well-established resting-state networks ^9–11^ (**Fig. S1-3**, see Supplementary results). Furthermore, we show that these states are organized in a hierarchical fashion, similar to what has been previously observed in humans ^7^. Together, these results show that spontaneous brain network activity in mice is characterized by similar principles of temporal organization as in humans, thus providing further evidence of how the mouse brain is a relevant model to causally study neuronal mechanisms of mammalian brain network dynamics.

### Single-area evoked neuronal spiking is sufficient to cause spatially distinct, fast brain network dynamics

What neural coding mechanism may support selective communication via oscillations? Selective communication relies on widespread, long-range anatomical connections among both cortical and subcortical networks. In a *divergent* case, where a single population of neurons sends signals to different projection targets (as opposed to a convergent case where multiple inputs converge to a single target), selective communication may arise through independent control of the gain for each of the projections targets. This would require *multiplexing* ^4^: the same spatiotemporal pattern of spiking in the projecting network contains separate information streams (or channels) that can be accessed by the different projection targets. In other words, different projection targets would be able to read out different signals from the same pattern of activity. If the multiplexing hypothesis in a divergent scenario were correct, one would predict that inducing activity with different neuronal spiking rates on top of a single-brain area basal activity would cause multiple patterns of circuit dynamics to emerge and to preferentially respond at different modulations. However, the hypothesis of multiplexing in a divergent scenario has not yet been causally tested.

Given its brain-wide projections and preferential modulation via the theta frequency band (5-10 Hz) ^12,13^, the entorhinal (EC)-hippocampal circuit represents a unique system to causally test the multiplexing hypothesis and thus understand brain network communication. Functionally, theta rhythms are proposed to have a key role in memory ^12,14^: it has been proposed that encoding the representation of novel memories, that relies on strong theta spiking input from the EC to CA1, is followed by a retrieval phase where strong communication from CA3 to CA1 allows to recall previously stored associations ^4^. However, it remains unknown whether selective information channels within the theta band allow the EC to access distinct downstream brain-wide projections, i.e. whether EC activity within the theta band employs multiplexing.

Understanding how brain function emerges from the synchronized firing of neurons, and whether multiple, distinct network dynamics can emerge from it, remain some of the most challenging questions in modern neuroscience. Answering these questions is fundamental to shed new light on circuit-specific mechanisms of information communication. Here, we combined HMM with optogenetics-evoked fMRI (ofMRI) ^15^ in mice with *channelrhodopsin-2* (ChR2)-transfected into excitatory neurons of the EC ^16^. Photostimulation was conducted with frequencies of 5, 10, and 20 Hz, pseudo-randomized between runs. Transfected EC neurons faithfully responded with frequency-locked action potentials to both extremes of the frequency-range tested(**Fig. S4**) (see Supplementary methods).

We asked whether localized excitatory neuronal induced-activity was sufficient to cause multiple, transient states of large-scale hemodynamic activity to emerge. Using data from N = 31 mice (N_WT-mCherry_ = 9, N_WT-ChR2_ = 10, N_3xTgAD-ChR2_ = 12), n = 153 runs, we sought to describe brain hemodynamics as a collection of 14 dynamic states (which yielded greater model similarity and lower free energy compared to other numbers of states; **Fig. S5a**). Each state captures a unique pattern of both amplitude and functional connectivity across brain regions (**Fig. S5, S6**). This 14-state HMM was then used to compare differences in brain network responses to optogenetic stimulation in the ChR2 and control groups (N_WT-ChR2_ = 10 mice; n = 54 runs; each animal underwent 6 runs acquired in 2 sessions; mCherry; N_WT-mCherry_ = 9 mice, n = 27 runs; each animal underwent 3 runs acquired in 1 session; all stimulation frequencies combined). We observed that the HMM was able to characterize changes in fast brain dynamics in response to the optogenetic stimulation in ChR2 animals (**Fig. 2a**, and with different states numbers as shown in **Fig. S7**) that was time-locked to the stimulation (**Fig. 2b**). Note that the HMM used here is an unsupervised method with no a-priori knowledge of the stimulation paradigm. As a comparison, a standard approach relying on mass-univariate GLM testing yielded one, single pattern of spatial activity in response to the optogenetic stimulation **Fig. S8**).

**Figure 2.**
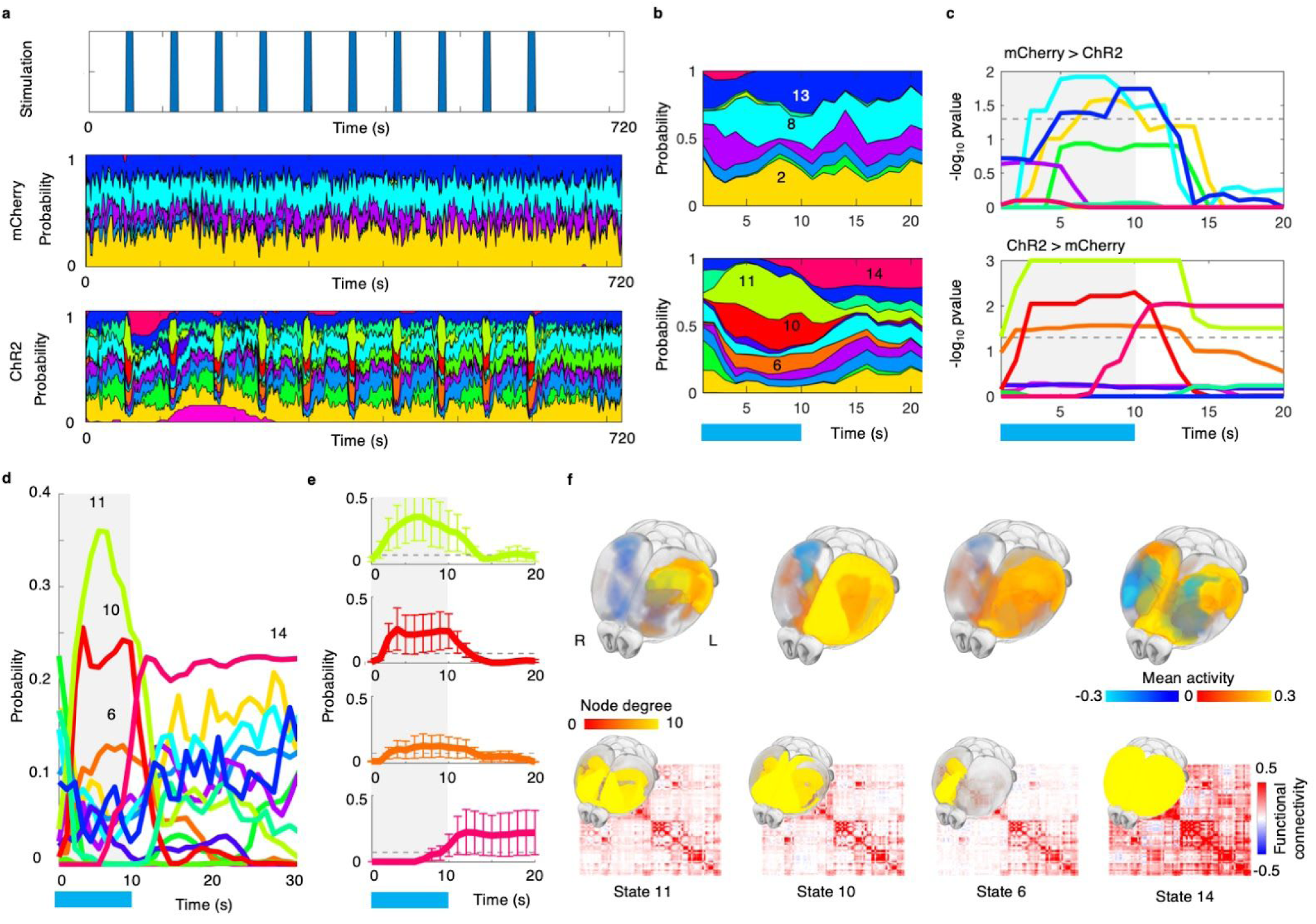
Optogenetic stimulation of the EC causes distinct states of brain network dynamics. A 14-state HMM was used to study brain network dynamics in response to EC stimulation at 5, 10, and 20 Hz in 3 groups of mice: N = 31 mice (N_WT-mCherry_ = 9, N_WT-ChR2_ = 10, N_3xTgAD-ChR2_ = 12), n = 153 runs (see **Fig. S5** for details on HMM fit). Here, however, only group comparison results in WT mice between mCherry controls and ChR2 groups is shown. **a**) (Top row) Optogenetic stimulation blocks shown in blue over time. (Middle and bottom rows) Group averages of HMM states activation probability aligned with optogenetic stimulation blocks. Each color represents a different HMM state. Middle row: WT-mCherry control group (N_WT-mCherry_ = 9 mice, n = 27 runs; each animal underwent 3 runs acquired in 1 session, all stimulation frequencies combined); bottom row: WT-ChR2 group (N_WT-ChR2_ = 10 mice; n = 54 runs; each animal underwent 6 runs acquired in 2 sessions, all stimulation frequencies combined). **b**) Area plot of group average responses for each state (colors as for **a**) over 20 seconds, time-locked to the optogenetic stimulation (shown by blue bar underneath) for control (top) and ChR2 group (bottom). **c**) Showing −log_10_ family wise error (FWE)-corrected p-values (across time and states) of group difference in HMM states activation probability time-locked to optogenetic stimulation (shown by blue bar and grey background). The threshold for statistical significance is indicated by the dotted line. **d**) HMM activation probability time-locked to stimulation shown in b) is here plotted as activation lines for ChR2 group only. **e**) Variability in HMM activation probability across subjects and runs (errors indicate standard deviation across subjects and runs in ChR2 group only; dotted line indicates the average activation level across all HMM states). **f**) brain maps of the 4 HMM states found to be significantly more active in the ChR2 group.

We then looked at the evoked brain network response to time-locked optogenetic stimulation. We focused on the HMM brain state activation probability at each time point, thus characterizing fast-transitions in brain network dynamics at a second-by-second resolution. Across blocks, for each time-point, we tested whether the probability of each HMM state being active was greater in the group expressing ChR2 compared to the mCherry controls (and vice versa). During stimulation, we found that 3 states (states 11, 10, and 6) were significantly more active in the ChR2 group, compared to controls (while correcting across time and states, **Fig. 2c**). Notably the spatial pattern of activity of these 3 states was significantly associated with the pattern of monosynaptic projection of EC, suggesting that these spatial patterns are direct results of inducing activation in excitatory neurons in EC (**Fig. S9a**). We also report that 3 states were significantly more active during stimulation in the control (mCherry) group (**Fig. S9b**). Their spatial pattern of activity, however, was not significantly associated with the pattern of monosynaptic projection of EC.

All these 3 states significantly respond to optogenetic stimulation showing (in the short-scale as opposed to the long-scale, see below) overlapping temporal responses (**Fig. 2d, e**). With a delayed onset, rise-time of around 5-7 seconds, and slower decay, the HMM-ofMRI evoked responses matched the dynamics of conventional stimulus-evoked BOLD fMRI ^17,18^.

Our HMM approach allowed us to characterize brain network states as dynamics in both BOLD amplitude fluctuations and functional connectivity (**Fig. S5, S6**). State 11 (**Fig. 2f**) shows greater BOLD amplitude (compared to the average across states) in the left EC (medial and lateral), subiculum, dentate gyrus, and CA1, and greater coupling bilaterally with the frontal cortex. State 10 shows a similar configuration of bilateral coupling but different BOLD amplitude: it shows greater activity in CA1, dentate gyrus, subiculum, orbital area, and in medial EC. State 6 shows greater activity in subiculum, dentate gyrus, medial EC, and CA1 and, of interest, left accumbens, pallidum, and thalamus, together with greater contralateral coupling in the frontal cortex. Together, these findings provide forward, causal evidence that inducing activation of excitatory neurons is sufficient to drive distinct brain network states of coordinated dynamics.

Of interest, we were also able to show that, in ChR2-transfected animals, state 14 showed significantly greater activation *after* the end of stimulation: this state was significantly present in ChR2-transfected animals but not in controls. State 14 showed greater BOLD amplitude bilaterally in the median EC, CA1, subiculum and amygdalar nuclei and, of interest, greater coupling across the whole cortex. This result suggests that after activation of excitatory neurons via optogenetic stimulation the brain does not necessarily return to a baseline or default mode state. Previous work, studying the optogenetic activation of the motor cortex, reported robust, delayed hemodynamic (and electrical) responses in the thalamus, an area with known axonal projections from the motor cortex ^18^. The authors attributed such response to network delays and interpreted it as an example of downstream activity ^18,19^. Downstream, delayed non-local activity (i.e. thalamic activity), was inferred to more likely depend upon a cascade effect that involves neuronal as well as non-neuronal elements, giving rise to more complex contributions to the vascular response ^20^. It is possible that we here observed how induced-neuronal spiking in genetically-targeted cells is sufficient to trigger a series of cascade effects, resulting in brain network entrainment into a non-default state, represented by state 14.

### Repetitive, single-area neuronal spiking is sufficient to cause long-scale causal interactions between brain network dynamics

As well as highlighting short-scale, time-locked responses to inducing neuronal activity, combining HMM with ofMRI also allowed us to characterize *long-scale* variations in responses. In other words, while standard ofMRI modeling requires averaging across multiple blocks in order to derive a single time-locked response, HMM also provided insight into how brain state responses to individual stimulation blocks varied over the course of the entire session (**Fig. 3a**). Here we compared ChR2 and control groups (N_WT-ChR2_ = 10 mice; n = 54 runs; each animal underwent 6 runs acquired in 2 sessions; mCherry; N_WT-mCherry_ = 9 mice, n = 27 runs; each animal underwent 3 runs acquired in 1 session; all stimulation frequencies combined). We first observed that while the long-scale pattern of stimulation response of state 11 remained constant throughout the whole stimulation session (**Fig. 3b**), state 10 showed a “U pattern” of response, with an initial decrease followed by a later increase in responsiveness (assessed through statistical comparison with mCherry group, See Supplementary methods). Of interest, we then observed that state 6 started responding more predominantly during the second half of the stimulation session, perhaps suggesting a long-scale process of circuit summation or amplification. As neuronal assemblies dynamically bind together in response to oscillatory synchronization, it is possible that the same augmenting property of resonators-oscillators takes place at the network level, thus providing a network-mechanism for the amplification of weak signals ^8,21^. At the small scale level (across and within cortical layers), strong axon collaterals, interacting in a recurrent excitatory feedback loop ^22^, are controlled by a variety of GABAergic interneurons ^23^, giving rise to a recurrent ensemble of excitation and inhibition. It is possible that this complex, small-scale interplay of feedback and feedforward effects, can give rise to large scale circuit amplification and inhibition that unfold dynamically over time.

**Figure 3.**
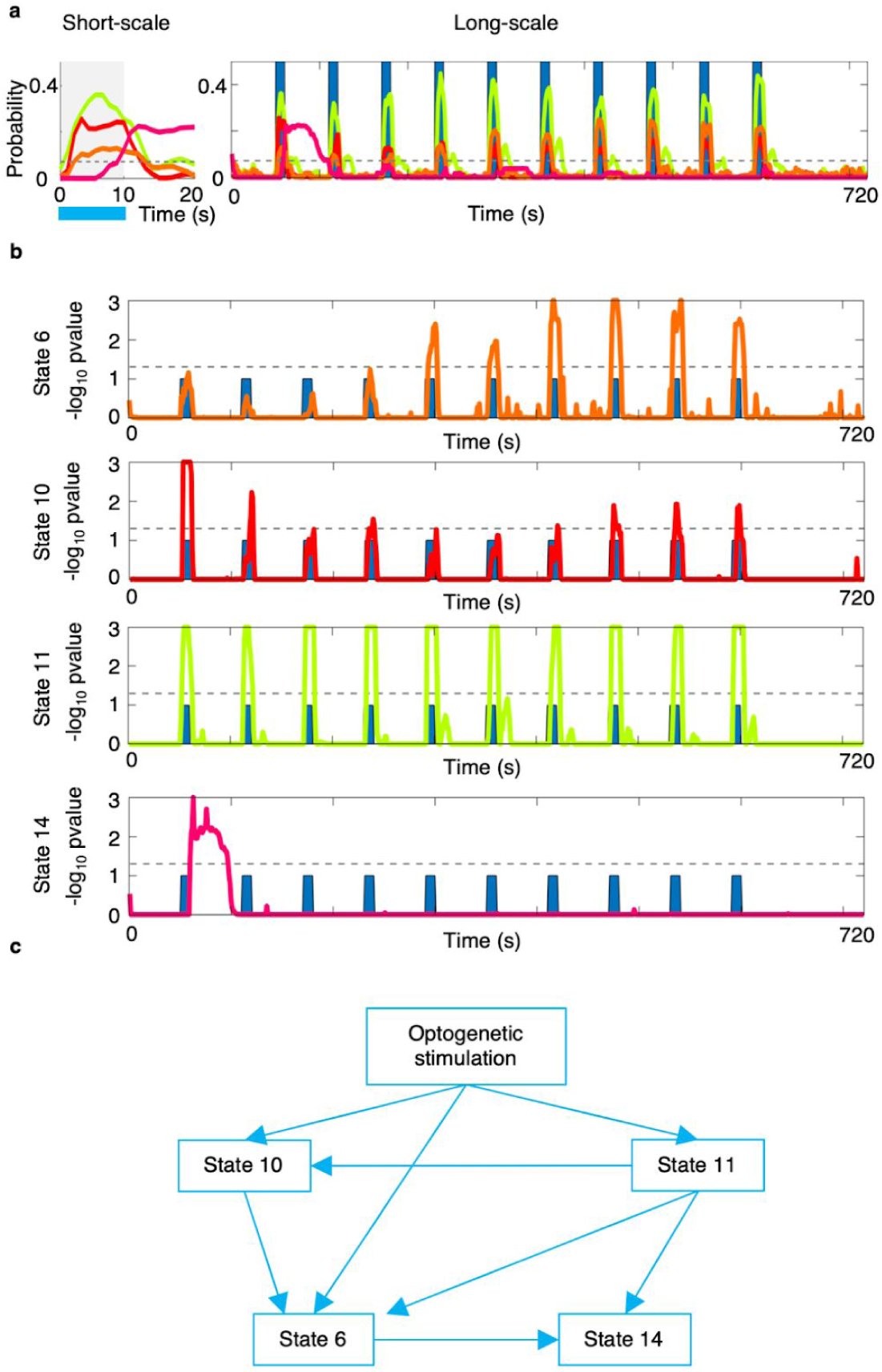
Optogenetic stimulation of EC causes long-scale interactions in brain network dynamics. HMM allows to characterize fast-scale (or time-locked) as well as long-scale responses (or slow variations across fMRI run) in whole brain hemodynamic activity. Results from analyzing ChR2 and control groups (N_WT-ChR2_ = 10 mice; n = 54 runs; each animal underwent 6 runs acquired in 2 sessions; mCherry; N_WT-mCherry_ = 9 mice, n = 27 runs; each animal underwent 3 runs acquired in 1 session; all stimulation frequencies combined). **a**) For WT-ChR2 group only, showing group average HMM states activation probability time-locked to stimulation (left) and variations across stimulation blocks (right). Dotted line indicates the average activation level across all HMM states. **b**) Showing −log10 FWE-corrected p-values of group difference (WT-ChR2 > WT-mCherry) in HMM states activation probability across time. Dotted line indicates statistical threshold. Then, across animals in WT-ChR2 group only, time-series of HMM states responding to optogenetic stimulation (state 6, 10, 11, and 14) were concatenated and modelled as a directed acyclic graph (DAG) (see Supplementary methods). **c**) Showing graph representation of inferred optimal DAG: arrows indicated Bayesian directed causality between optogenetic stimulation and state activity.

Feedforward and feedback mechanisms modulating the unfolding of brain network dynamics over time may further imply directed interactions between states of activity. In order to formally test this prediction, across all WT-ChR2 mice, we modelled on/off optogenetic stimulation together with HMM states’ activation probability as a directed acyclic graph (DAG) (See Supplementary methods). We looked for the optimal Bayesian network allowing us to establish statistical conditional dependencies (edges) between variables (nodes; here on/off stimulation and HMM time courses for states 6, 10, 11, and 14) in order to estimate direct causal connections between stimulation and HMM states. In other words, this approach allowed us to determine causality between states’ time courses. The optimal causal Bayesian network analysis (See Supplementary methods) (BIC = 684,159) indicates the following chain of events (**Fig. 3c**): optogenetic stimulation directly drives activation of state 11, state 10, and state 6; state 10 however, is also influenced by onset of state 11; and state 6 is also influenced by onset of states 11 and 10. Crucially, the same Bayesian network also showed that activation of state 14 is not directly caused by optogenetic stimulation; however the stimulation effect is mediated through the activation of states 11, 10, and 6. This DAG provides causal, directional evidence that single-area neuronal spiking is sufficient to drive multiple brain network dynamics and their non-trivial interactions, further showing how inducing neuronal excitation may result in an activity propagation or cascade effect, possibly modulated by the interplay of circuit-level feedforward and feedback interactions unfolding over time.

Monitoring the effects of brain stimulation is of crucial importance not only for basic research, but also for translational research aiming to predict patient responses to brain stimulation. Here, we show that dynamic states might be an effective, data-driven metric for such monitoring. In particular, differential effects between short-scale and long-scale brain stimulation protocols have been widely reported, not only for invasive techniques in rodents, like chemogenetics ^24,25^, but also for non-invasive brain stimulation techniques such as TMS ^26^ and tDCS ^27^ in humans. These differential effects are often visible not only in physiology ^28^, but also in behavioural effects ^24,25^, and understanding them is key for further translation into clinical populations. Here, imaging temporal dynamics together with brain stimulation uncovers one potential mechanism underlying such timescale-driven effects: changes in network dynamics. These effects may provide a parsimonious explanation for the constellation of previously reported timescale-driven effects of brain stimulation, and a mechanism for future studies to consider when designing and testing non-invasive brain stimulation paradigms. Given the ubiquity of timescale-dependent phenomena in brain stimulation, these results generate hypotheses that will be easily testable in a number of different non-invasive brain stimulation paradigms across basic and clinical contexts.

### Distinct brain network dynamics preferentially respond to different stimulation frequencies within the theta frequency band

A large body of literature indicates how human brain functional specialization relies on the transient synchronization of oscillations ^5,29^. As oscillatory dynamics emerge from the interplay between cellular and circuit mechanisms, these same resonant-oscillatory features are thought to allow neuronal networks to select inputs based on their frequency characteristics (for a review: ^8^). Because of this resonant-oscillatory property, the hypothesis of oscillatory multiplexing for selective communication would predict that neuronal assemblies (or dynamic brain networks) have developed an input frequency preference (or gain-response function) ^4^, thus allowing for flexible responses to input frequency changes.

Neuronal oscillations based on theta modulation represent the basis of EC-hippocampal circuit communication and are thought to support memory encoding and recall ^12,13^. Here, we first asked whether distinct, optogenetically-driven brain network dynamics show preferential activation as a function of neuronal firing rate. To answer this question, in the WT-ChR2 group (N_WT-ChR2_ = 10 mice; n = 54 runs; each animal underwent 2 sessions of 3 runs, one run per stimulation frequency (5, 10, 20 Hz)), we focussed on the HMM states which showed time-locked increases in response to stimulation (states 6, 10, 11, and 14), and assessed their response to stimulation at 5 Hz (compared to 10 and 20 Hz) and at 10 Hz (compared to 5 and 20 Hz). Using a repeated measure ANOVA (see Supplementary methods), we tested the effect of frequency on HMM state probability onset. We found a significant overall (F-test) effect of inducing neuronal firing with theta frequency band stimulation on state 10 and state 11 (**Fig. 4a**). Crucially, the post-hoc analyses (T-tests) showed that while state 10 responded significantly more at 10 Hz stimulation (compared to 5 and 20 Hz), state 11 responded more at 5 Hz stimulation (compared to 10 and 20 Hz) (**Fig. 4b, S10b, c**). To test whether these frequency-specific effects were the result of a greater EC responsivity as a function of increasing stimulation frequency (5 Hz < 10 Hz < 20 Hz), using a repeated measure ANOVA we tested for a monotonic (either linear or nonlinear) increase in HMM states’ response. No significant result was found. As a further, post hoc control test, we tested for a specific response to 20 Hz stimulation (compared to 5 and 10 Hz). No significant result was found. These results provide forward evidence for multiplexing as a mechanism for flexible communication in brain networks, causally linking theta modulation in excitatory neurons with distinct spatiotemporal patterns of brain network dynamics.

**Figure 4.**
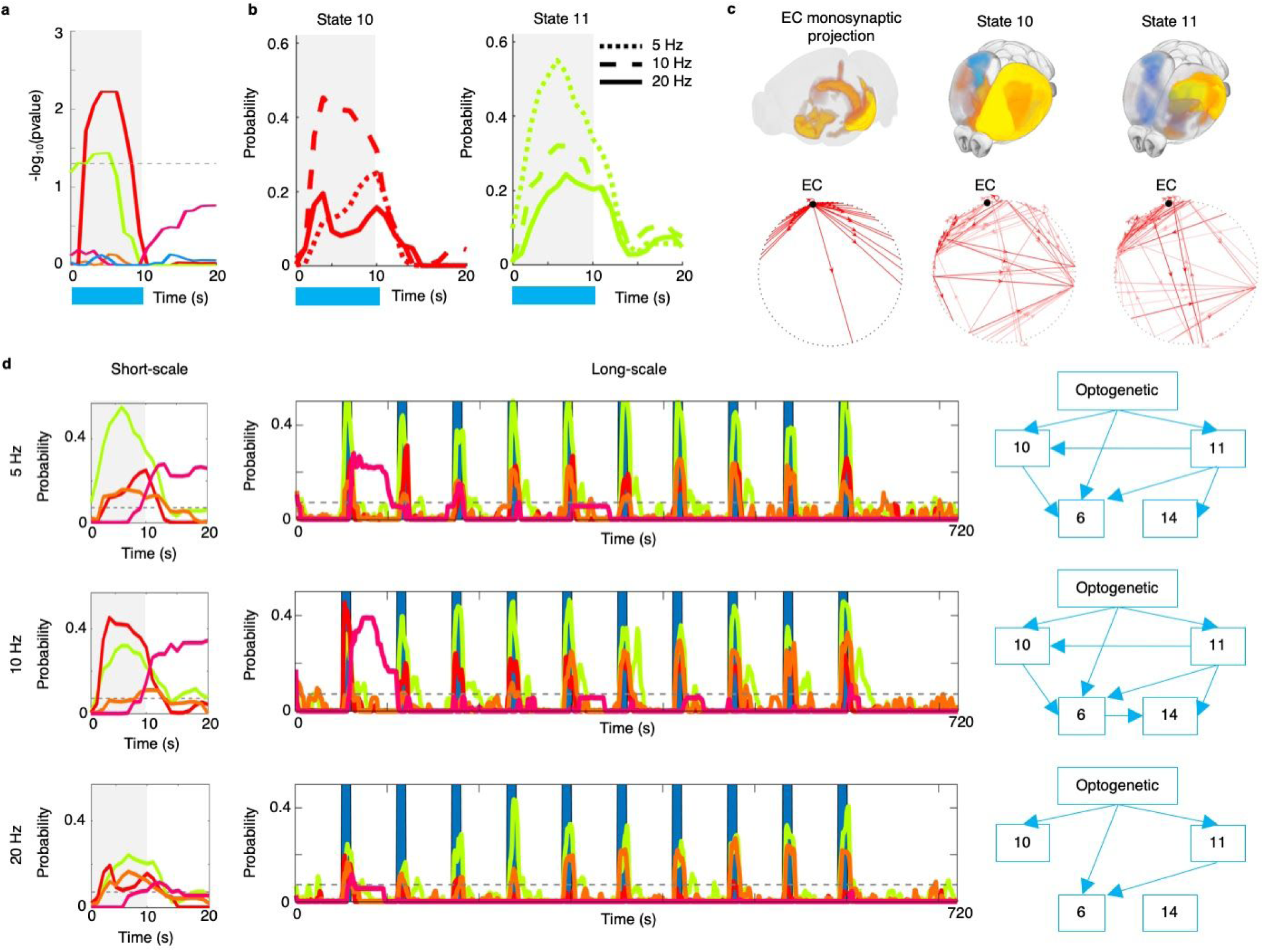
Varying frequency of EC neuronal spiking entrains the brain into distinct states of network dynamics. Results from analyzing only the WT-ChR2 group (N_WT-ChR2_ = 10 mice; n = 54 runs; each animal underwent 2 sessions of 3 runs, one run per stimulation frequency (5, 10, 20 Hz)). Stimulation frequencies were explicitly modelled in the statistical tests. The multiplexing hypothesis predicts that in a divergent scenario multiple target regions would respond differently to the same stimulation input. A repeated measure ANOVA was used in order to test the effect of varying stimulation frequency on HMM states activation probability. **a**) Showing −log_10_ FWE-corrected p-values for an overall effect of frequency on HMM states activation probability (F-test results). Dotted line indicates statistical threshold. **b**) HMM activation probability for states 10 and 11 during stimulation at 5, 10, and 20 Hz (See **Fig. S8** for results from T-test). **c**) Showing EC monosynaptic projections as well as predicted anatomical circuits underlying the activity pattern of states 10 and 11(See **Fig. S10, S11** for details). The effect of varying stimulation frequency on long-scale brain network dynamics was then tested. For each stimulation frequency (5, 10, 20 Hz), HMM states time-series across animals were concatenated and a DAG was inferred. D) Respectively for each stimulation frequency, showing short-scale (left), long-scale (middle) HMM states dynamics, and DAGs of directed causal interactions between stimulation and HMM states (right).

If inducing activation with given firing rates on top of basal activity in a single population of neurons results in distinct brain network dynamics, it follows that the same signal must rely on independent gain control at the level of the target regions ^4^. As one previous study has shown that different patterns of (static) activity may rely on different sets of anatomical connections ^30^, one hypothesis could be that gain control in response to varying stimulation frequency is implemented in the form of different anatomical circuits. If this were the case, it would provide an anatomical basis for distinct information channels. We therefore tested the hypothesis that states preferentially responding at different frequencies do rely on specific anatomical circuits. To test this hypothesis we performed structural-to-functional mapping (see Supplementary methods, **Fig. S11** for explanatory cartoon), characterizing the underlying patterns of white-matter anatomical projections (or circuit) best predicting functional activity in brain network dynamics. Using the connectivity matrix of directed tracing projections publicly available from the Allen Brain Institute ^31^, we entered all areas’ patterns of outgoing anatomical projections (each column is a brain area, for a single column each row represents the amount of outgoing projections to a target area) in a leave-one-out cross-validated Ridge regression in order to predict the activity pattern of states 10 and 11 (rows are BOLD amplitude values of, respectively, HMM state 10 and, independently, 11). This yields significant predictions of HMM states activity patterns of HMM states 10 and 11 (**Fig. S10c**: state 10: predicted rho = 0.82 (R^2^ = 0.67), cross-validated deviance = 29.66, p-value = 0.001; **Fig. S10h**: state 11: predicted rho = 0.77 (R^2^ = 0.60), cross-validated deviance = 35.72, p-value = 0.001; significance assessed with 1,000 permutations). We then extracted the average regression coefficients across folds, and multiplied these values by the outgoing pattern of projections in order to show the anatomical circuits underlying activity states 10 and 11 (**Fig. S10e, l**). We then assessed the specificity of these anatomical circuits in predicting the opposite state of activity (i.e. regression betas estimated in prediction of state 11, now used to predict state 10 activity, and vice versa). We found that using the regression coefficients of state 11 to predict state 10 activity yielded a statistically significantly worse prediction (n = 90 brain parcels; see **Table S1**; comparing correlations from dependent samples using Fisher’s Z transformation for correlation coefficients: Z = 2.472, p-value = 0.007) and, vice versa, using regression coefficient of state 10 to predict state 11 activity also let to a significantly worse prediction (n = 90; see **Table S2**; comparing correlations from dependent samples: Z = 2.296, p-value = 0.011). Together, these results show that the two directed structural circuits, despite sharing a portion of connections linked to the EC, are indeed different and specifically relate to one but not the other state of activity, suggesting a role for network topology in multiplexing: different anatomical circuits may provide an anatomical basis for distinct information channels.

We next asked the question of whether inducing oscillations in a single area at different frequencies drives long-scale brain network dynamics into different trajectories of activity. For each frequency of stimulation (5, 10, and 20 Hz), we considered the long-scale time-series of the HMM states previously identified and built three, separate DAGs representing the directed causal influences between brain dynamics and optogenetic stimulation. We observed that although DAGs in the theta frequency band (5 and 10Hz) were highly overlapping, inducing neural firing in the beta band (20 Hz) drove the system into a distinct configuration with far less interactions between brain dynamics (**Fig. 4d**) and where the downstream activity state (state 14) was not accessed (BIC values > 210,000 for all frequency conditions) (**Fig. S12**, see Supplementary results). These findings crucially show how inducing activity in the EC at different frequency bands is causally sufficient to drive the brain network into trajectories that unfold differently over time, thus probing different long-scale, complex configurations of interactions between brain areas: a necessary condition for flexible computation.

### Impaired multiplexing in brain network dynamics in a transgenic model of Alzheimer’s disease

Alzheimer’s Disease (AD) is a progressive neurodegenerative disorder affecting the entorhinal cortex and the hippocampus. AD is caused by neuronal degeneration, leading to brain-wide dysfunction and cognitive deficits ^32,33^, but both the cell-level mechanisms and network-level mechanisms ^34^ that unfold during AD are poorly understood. A step forward has been the identification of familial, early onset forms of the disease, which allowed the generation of animal models, and thus the exploration of the potential pathophysiological mechanisms behind the disorder ^35,36^.

Here, we used a well-established model of familial AD to recapitulate behavior-relevant ^37^, naturalistic dysfunction of EC circuits, and probe how such dysfunction may affect network dynamics and multiplexing in the nervous system. Using ofMRI, we found that 3xTgAD mice are characterized by an altered configuration of baseline brain activity (state 4) (**Fig. 5a-d**), and that theta-rhythms stimulation resulted in a reduction of its activation probability that outlasted the duration of stimulation (**Fig. S13**) (see Supplementary results). These results show that AD mice, despite showing aberrant brain dynamics (state 4) at baseline, may return to wild-type-like brain dynamics following theta rhythm optogenetic stimulation of EC. This complements previous evidence that EC stimulation may provide therapeutic benefit in AD ^38^, and provides causal evidence of a mechanism that might underpin clinical benefits of EC stimulation.

**Figure 5.**
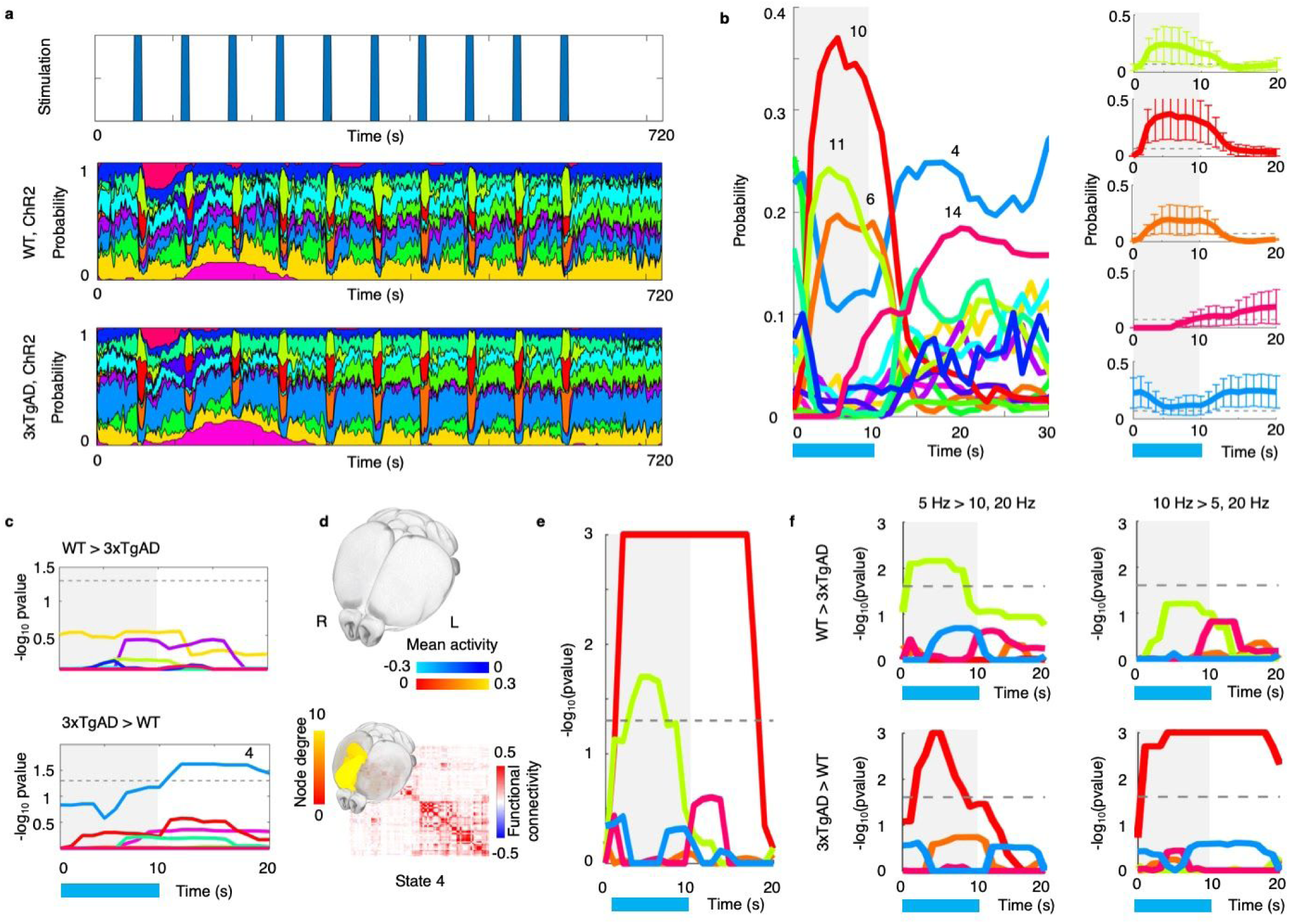
Alzheimer’s disease impairs the preferential HMM states responses to varying spiking frequency in EC. Results of group comparison between WT-ChR2 group and 3xTgAD-ChR2 group. **a**) Group average HMM activation probability aligned with optogenetic stimulation blocks, respectively for WT-ChR2 group and 3xTgAD-ChR2 group (N_WT-ChR2_ = 10 mice; n = 54 run; each animal underwent 6 runs acquired in 2 sessions; N_3xTgAD-ChR2_ = 12 mice; n = 72 runs, each animal underwent 6 runs acquired in 2 sessions; all stimulation frequencies combined). **b**) For 3xTgAD-ChR2 group only, average HMM response time-locked to stimulation and how it varies across mice and runs. **c**) Showing −log_10_ FWE-corrected p-values (across time and states) of group difference in HMM states activation probability. Dotted line indicates statistical threshold. **d**) Brain map of “aberrant activity” (state 4) present in AD. Mean activity showed low absolute values in comparison to other HMM states. We then studied whether multiplexing is impaired in AD. Here, stimulation frequencies were explicitly modelled in a mixed-design ANOVA (see **Fig. S14a** for design matrix). Statistical analyses were carried out on WT-ChR2 group and 3xTgAD-ChR2 group (N_WT-ChR2_ = 10 mice; n = 54 run; N_3xTgAD-ChR2_ = 12 mice; n = 72 runs; for both groups, each animal underwent 2 sessions of 3 runs, one run per stimulation frequency (5, 10, 20 Hz)). **e**) Showing −log_10_ FWE-corrected p-values for an overall interaction effect of group *by* (theta-) frequency on HMM states activation probability (F-test results). **f**) Showing −log_10_ FWE-corrected p-values for post-hoc T-tests of interaction analysis. Dotted line in **c, e, f**, indicates statistical threshold.

Then, building on the fact that distinct states of brain network dynamics preferentially respond to different EC stimulation frequencies in wild-type mice, we tested whether this frequency-specific response was different between AD and WT mice. We found that there was a significant interaction effect of group by theta-frequencies on state 10 and 11 (**Fig. 5e, S14a, b**) (see Supplementary results). Specifically, while WT mice showed state 10 to preferentially respond at 10 Hz stimulation and state 11 at 5 Hz, in AD mice state 10 did not show a preferential responding at *both* 5 and 10 Hz while state 11 did not show any significant effect of variation in neuronal spiking frequency (**Fig. 5f, S14c-f**). These results provide causal evidence that 3xTgAD mice lack differential brain network response to varying EC neuronal spiking rate, thus showing how neural coding for flexible brain network communication in AD mice is impaired.

## Discussion

Brain network communication emerges as an organic property from the interactions between billions of cells linked together in interconnected circuits. Although oscillations in neuronal populations are thought to represent the medium for information transfer ^8^, the neural coding that allows for flexible, selective communication to emerge has been unknown.

In this work, we considered the EC, a well-known hub in the EC-hippocampal memory system known to rely on theta rhythms to mediate memory encoding and recall ^12^. We used this structure and its anatomical projections to causally test the communication mechanism of multiplexing in a divergent networked scenario, in health and in AD.

We provide sufficient causal evidence that, in healthy WT mice, driving theta oscillations in EC neuronal spiking causes distinct brain network dynamics to emerge (**Fig. 2**), with precise short- and long-scale dynamics (**Fig. 3**). Crucially, we then provide evidence that within the same theta band, spatially distinct but temporally overlapping networks preferentially respond to inducing oscillations at either 5 or 10 Hz (**Fig. 4**), revealing how multiple information streams within the same theta band originate from a common neural substrate using frequency-division multiplexing. Computationally this is achievable because the gain modulation reached at one specific frequency is independent from the average gain experienced at other frequencies ^4^. Furthermore, we show that these distinct functional information streams can rely on different anatomical circuits (**Fig. 4, S10**), thus providing evidence of a role for topology in network communication. As the EC-hippocampal memory system is known to rely on theta oscillations as a medium for neuronal communication ^12^, this work provides a causal forward test for oscillation multiplexing as a neural code sufficient for achieving flexible information routing in brain networks.

Notably, we also show that in a transgenic model of AD, the EC mechanism of neuronal multiplexing is impaired: we observe the lack of distinct information streams within theta rhythms (**Fig. 5, S14**), thus impairing flexibility in brain network communication. This result provides mechanistic causal evidence of how the early stages of AD neurodegeneration may cause an alteration in the neural code used for flexible communication in brain networks, thus providing novel insight into the aberrant mechanisms underlying cognitive decline.

The EC has long been implicated in memory formation and spatial navigation ^39^, with recent studies highlighting a potential role for cognitive maps in reconciling this apparent dichotomy in the function of EC ^40^. Previous studies deploying optogenetics to EC have shown that inhibition of cells in the EC has downstream effects on the hippocampus, impairing memory formation in fear-conditioning paradigms ^41,42^. Future studies should establish the role of multiplexing in memory formation and in navigation of abstract spaces.

Previous work has explored how optogenetic stimulation at different frequencies may also result in different magnitudes of regional activation and static functional connectivity ^43–45^. Furthermore, a recent study has also demonstrated how driving neuronal activity with optogenetics results in static patterns of effective connectivity as determined through dynamical causal modeling ^46^. Here, we employ a novel computational approach and provide evidence that driving neuronal spiking through means of optogenetics results in network states of dynamic brain activation and dynamic functional connectivity, across short- and long-time scales, rather than exhibiting one single, static, pattern across time. Capturing spatio-temporal dynamics of brain activity also allows us to use ofMRI to pose more fundamental questions about neural coding. For example, building on previous evidence and known concepts of neuronal oscillatory synchronization ^8^, we also provide causal evidence that varying neuronal spiking rate in the same brain region entrains the brain into spatially and temporally distinct network states that show frequency response preferences, providing novel mechanistic understanding of how selective information streams may arise from multiplexed neuronal activity.

As with any methodology, ofMRI also presents limitations. Here, we used optogenetics to probe excitatory neuronal spiking in the EC and observed short-scale and long-scale brain network dynamics through means of whole brain BOLD fMRI. However, despite using HMM in order to access fast transitions in BOLD fluctuations amplitude and coupling, brain imaging techniques have an intrinsic trade-off between a second-resolved temporal resolution and whole brain imaging. We cannot ignore therefore the fact that perturbing genetically-targeted excitatory neurons inevitably also led to an immediate engagement of local perisynaptic activity in the form of the excitatory and inhibitory loops mentioned above ^20^. However, implementing HMMs ^57^ allows us to disentangle dynamics of spatially and frequency distinct networks otherwise temporally overlapping in the short-scale and not observable using standard analyses or invasive techniques like single-area electrophysiology. Furthermore, using HMMs, together with experimentally varying stimulation frequency, allowed us to characterize between network states with a long-scale response amplification and network states preferentially responding at specific theta frequencies – suggesting that different, underlying neural mechanisms are taking place.

Our experimental and computational approaches, therefore, allowed us to establish the causal *sufficiency* ^47^ of oscillatory synchrony as a mechanism for flexible communication in brain networks. Future research should leverage optogenetic *inactivation* in order to test the causal *necessity* of cell-type-specific spiking, and to characterize the specific role of other neural populations.

To truly understand how the brain functions, one must ultimately understand causal cell-type-specific mechanisms supporting selective brain network communication. Combining optogenetics with concurrent fMRI allows causal network neuroscience: causally interfering with neuronal dynamics to understand brain network design. However, because within each brain circuit there is considerable sub-circuit specificity, intermingled neurons with different patterns of long-range projections can modulate circuit function in remarkably selective ways ^47^, thus probing distinct information streams in brain network function. In this work, modulating the firing frequency of a single-area specific cell-type has allowed us to test the multiplexing hypothesis in a divergent networked scenario: differential oscillatory synchrony is a *sufficient* neural code to allow for selective and flexible communication in brain network dynamics.

## Acknowledgements

This work was supported by the Singapore BioImaging Consortium awards to JG (#2016) and FM (#2017). HJB is supported by a Wellcome Principal Research Fellowship (110027/Z/15/Z). The Wellcome Centre for Integrative Neuroimaging is supported by core funding from the Wellcome Trust (203139/Z/16/Z). We would also like to thank Anderson Winkler for his helpful comments on statistical testing.

## Author contributions

PS, AL, and JG designed the research; FM acquired the data; PS analyzed functional MRI data; PS wrote the manuscript. All authors contributed with the interpretation of the findings and in writing the manuscript.

## Supplementary information

### Supplementary results

#### Mouse brain states of spontaneous activity are hierarchically organized in time

In the rs-fmri dataset, spontaneous whole-brain BOLD fMRI fluctuations from 29 anesthetized mice (N_WT_ = 10; N_3xTgAD_ = 19; for a total of n = 52 runs), were modelled through a 14-state HMM state (**Fig. S1**). This number was chosen in order to be consistent with the primary analysis of concurrent ofMRI. We show that, as in humans, rodent brain states transitions are not completely at random, although in a stochastic fashion. This is represented by the state-to-state transition probability matrix (**Fig. S1a**) and results in spontaneous, fast transitioning on a second-to-second scale (**Fig. S1b**). The mean amplitude and functional connectivity maps for all states are shown in **Fig. S1c, d, S2**. We then studied whether, similarly to humans ^48^, the mouse brain displays a hierarchical organization of its temporal features. Across multiple HMMs with different numbers of states (10, 12, 14, 16 and 18) we calculated fractional occupancy (FO), the time an animal spent in one state over the course of the rs-fmri session. Across mice, we then created a meta-correlation matrix estimating similarities in FO across multiple HMMs (**Fig. S3a**). In a data-driven fashion, we then estimated the best number of metastates. On the meta-correlation matrix, we used a multi-scale community detection algorithm to characterize sets of states (or modules) with similar FO distributions across mice, also defined as metastates. We found that, across a range of values in the resolution parameter, maximum mean partition similarity was achieved for γ = 1 (**Fig. S3b**), hinting at the presence of 3 well-defined metastates. The resolution parameter (γ = 1) with higher partition similarity was then applied to the single resting-state fMRI HMM run with a fixed number of states (k = 14): this led to the same result of 3 well defined metastates (**Fig. S3c, d**). This allowed us to characterize a hierarchical temporal structure of rodent brain states that is remarkably similar to that of the human brain.

#### Causal role of theta-band neuronal spiking in causing downstream activity

We then studied the response of the downstream state of activity (state 14) involving greater activation bilaterally in the median EC, Ammon’s horn, subiculum and amygdalar nuclei. Although the time-locked analysis averaging across blocks showed a robust and significant response starting *after* optogenetic stimulation, the long-scale analysis revealed that such response took place in a statistically significant and long-lasting manner only after the first block of stimulation (**Fig. S12a**). This result would not otherwise have been observable with standard modeling approaches to ofMRI.

Here, we tested the relevance of theta rhythms in causing downstream, delayed activity. We tested whether driving brain network dynamics with theta oscillation (but not beta) could temporally precede and quantitatively explain the magnitude of downstream state activity. Across animals, we found that, when driving theta oscillations (5 and 10 Hz), inter-individual differences in state 10 activation probability *during* optogenetic stimulation (51-60 seconds from beginning of ofMRI run) were positively associated with differences in downstream activation *after* stimulation (61-110 seconds). Crucially, this was not the case when brain network dynamics were entrained with beta rhythms (20 Hz) (**Fig. S12b**, Spearman’s correlations were Bonferroni corrected across all active states and frequencies: Bonferroni corrected p-values were considered significant if < 0.0056).

These results, together with the fact that the EC projection pattern alone can not explain the spatial pattern of the downstream state of activity (**Fig. S9a**), argue in favour of a networked and frequency-band specific origin: the downstream, delayed activity state was the non-random result of previously driving theta oscillations and entraining a EC-hippocampal-orbitofrontal network (state 10), suggesting therefore a system-specific rebound mechanism.

#### Altered baseline activity and impaired multiplexing in brain network dynamics in a transgenic model of Alzheimer’s disease

Using ofMRI, we first tested how HMM states of brain network hemodynamic activity are affected in the 3xTgAD model of AD compared to wild-type mice (N_WT-ChR2_ = 10 mice; n = 54 runs; each animal underwent 6 runs acquired in 2 sessions; 3xTgAD-ChR2, N_3xTgAD-ChR2_ = 12 mice; n = 72 runs, each animal underwent 6 runs acquired into 2 sessions; all stimulation frequencies combined). We first asked whether activation probability of HMM states was different between ChR2 and 3xTgAD mice. While many aspects of optogenetic stimulation-induced activity were comparable (**Fig. 5a-d**), we found a significant, overall increase in activation probability *after* stimulation in 3xTgAD mice in a state characterized by greater connectivity in the right frontal cortex (state 4, Fig. 5d). However, we also observed in **Fig. 5a** that this higher level of activation is present throughout the entire ofMRI run, thus representing a baseline state. Optogenetic stimulation therefore does not result in an increase of state activation *after* stimulation but in a reduction activation probability *during* stimulation, suggesting that even at early onset stages, AD may be characterized by an altered configuration of dynamic activity. Furthermore, we also observed that the spatial pattern of functional activity of this aberrant state was positively associated with EC monosynaptic projections (**Fig. S9c**), suggesting some complex interplay between EC connectivity and response to EC stimulation in AD mice.

Having found an altered state of AD brain dynamics (state 4) compared to WT mice, and that its activation probability was decreased during each instance of optogenetic stimulation, we then asked whether its response underwent a group *by* theta-frequency reduction in AD mice (equivalent to group by beta-frequency increase in AD mice). Using a mixed-effect ANOVA (see Supplementary methods), we tested a group *by* 20 Hz interaction effect (N_WT-ChR2_ = 10 mice; n = 54 runs; N_3xTgAD-ChR2_ = 12 mice; n = 72 runs; for both groups, each animal underwent 2 sessions of 3 runs, one run per stimulation frequency (5, 10, 20 Hz)). We found that activation probability of the altered state was greater in AD compared to WT mice when stimulation was induced at 20 Hz compared to 5 and 10 Hz (**Fig. S13a**). In other words, theta rhythms (5 or 10 Hz) were more effective than beta rhythms (20 Hz) stimulation at reducing activation probability of the altered state of AD brain dynamics. Although state 4 had greater activation probability in AD-like mice compared to WT mice, it is still possible that in WT mice state 4 undergoes the same effect of theta frequency. The result of this group *by* frequency interaction however shows that it is not the case: WT mice do not show the same effect of frequency for state 4, implying that this frequency-specific effect is present only when state 4 is pathologically over-active (**Fig. S13a, S14b**). As we have shown that repetitive neuronal spiking is sufficient to induce long-scale dynamics (Fig. 3), we then asked whether reducing state 4 activation probability through means of theta rhythms also resulted in a long-scale effect. For state 4, across stimulation blocks, we calculated the subject specific average in activation probability *after* stimulation ended (as each block lasted 60 seconds, and stimulation was on for 10 seconds, we considered the time interval between 11 and 60 sec). Using a repeated measure ANOVA (see Supplementary methods) only on AD mice (N_3xTgAD-ChR2_ = 12 mice; n = 72 runs; each animal underwent 2 sessions of 3 runs, one run per stimulation frequency (5, 10, 20 Hz)), we tested the effect of varying neuronal spiking rate. We found an overall effect of theta frequencies on state 4 activation probability even after stimulation ended (F-test: p = 0.0100), resulting in a significant reduction of state 4 probability when neuronal firing was driven at 5 and 10 Hz compared to 20 Hz (post-hoc T-tests: respectively, 5 Hz stimulation compared to 10 and 20 Hz, p = 0.0130; and 10 Hz stimulation compared to 5 and 20 Hz, p = 0.0020) (**Fig. S13b**), showing that reductions in state 4 activation through means of theta rhythms outlast the duration of optogenetic stimulation. Taken together, these results show that AD mice, despite showing aberrant brain dynamics (state 4) at baseline, may return to WT-like brain dynamics following theta rhythm optogenetic stimulation of EC.

Then, building on the fact that distinct states of brain network dynamics preferentially respond to different EC stimulation frequencies in WT mice, we tested whether this frequency-specific response was different between AD and WT mice (N_WT-ChR2_ = 10 mice; n = 54 runs; N_3xTgAD-ChR2_ = 12 mice; n = 72 runs; for both groups, each animal underwent 2 sessions of 3 runs, one run per stimulation frequency (5, 10, 20 Hz)). Using a mixed-effect ANOVA (see Supplementary methods), we tested a group *by* (theta) frequency interaction on HMM state probability onset. We found a significant overall interaction effect (F-test) of group *by* neuronal firing frequency on state 10 and state 11 (**Fig. 5e, S14a, b**). The post-hoc interaction analyses (T-tests) showed that, while state 11 response at 5 Hz stimulation (compared to 10 and 20 Hz) was significantly greater in WT compared to AD mice, state 10 response was significantly greater in AD compared to WT mice at both 5 and 10 Hz stimulation (compared to 20 Hz) (**Fig. 5f**). These results show that brain network response to varying stimulation frequency significantly differed between groups.

Specifically, while WT mice show different dynamic brain networks to preferentially respond to varying stimulation frequency (state 10 to 10 Hz and state 11 to 5Hz), in AD mice state 10 showed a preferential response at *both* 5 and 10 Hz while state 11 did not show any significant effect of variation in neuronal spiking frequency. A further, separate repeated measure ANOVA only on AD mice confirmed this result (N_3xTgAD-ChR2_ = 12 mice; n = 72 runs; each animal underwent 2 sessions of 3 runs, one run per stimulation frequency (5, 10, 20 Hz)) (**Fig. S14c-f**). Together, these results provide causal evidence that 3xTgAD mice lack differential brain network response to varying EC neuronal spiking rate, thus showing how neural coding for flexible brain network communication is impaired in AD mice.

## Supplementary methods

### Sample

All experiments performed in Singapore Bioimaging Consortium, A*STAR, Singapore, were in accordance with the ethical standards of the Institutional Animal Care and Use Committee (A*STAR Biological Resource Centre, Singapore, IACUC #171203). Male 3xTgAD and WT mice on the same background strain (129sv/c57bl6) were used for the rs-fmri and the ofMRI experiments. The colonies of both 3xTgAD and WT mice were maintained ‘in-house’ through the pairing of homozygous individuals. Mice were housed in cages of up to five, with same-sex and genotype cage-mates in a pathogen-free environment, kept at a 45-65% humidity, under a 12:12-hour light-dark cycle and room temperature, with *ad-libitum* access to food and water.

Two datasets of animals have been used in this work: a rs-fmri dataset (Mouse_rest_3xTG, 10.18112/openneuro.ds001890.v1.0.1) and an ofMRI dataset (Mouse_opto_3xTG, 10.18112/openneuro.ds002134.v1.0.0). Wild-type and 3xTgAD mice underwent rs-fmri sessions at 3 and 6 months of age, longitudinally with the cryogenic coil setup. The second dataset included wild-type and 3xTgAD mice that under firstly underwent optogenetic surgery for CaMKIIa-mCherry or CaMKIIa-ChR2-mCherry injection (without opsin and with opsin, respectively) and, 3 weeks post injection, underwent the ofMRI protocol with the single-loop coil, with 5/10/20 Hz stimulation paradigm. Specifically, one control group was injected with CaMKIIa-mCherry (N_WT-mCherry_ = 9). One control group and one 3xTgAD group were injected with CaMKIIa-ChR2-mCherry (N_WT-ChR2_ = 10; N_3xTgAD-ChR2_ = 12, respectively) and were longitudinally scanned also at 6 months of age (N_WT-ChR2_ = 8; N_3xTgAD-ChR2_ = 10). For the specific breakdown of the number of animals and runs per group, see **Table S3**.

### Viral injection

Optogenetic surgery procedure is described in ^49^. Briefly, WT and 3xTgAD mice (∼30 g, ofMRI dataset: N = 31 mice (N_WT-mCherry_ = 9, N_WT-ChR2_ = 10, N_3xTgAD-ChR2_ = 12) were anaesthetised with a mixture of ketamine/xylazine (ketamine 75 mg/kg, xylazine 10 mg/kg). The head was shaved and cleaned with three wipes of Betadine® and ethanol (70%). Lidocaine was administered subcutaneously, *in situ*. To avoid hypothermia, animals were placed on a warm pad and the head fixated on the stereotaxic frame; protective ophthalmic gel was applied to avoid dryness. After removing the scalp, a craniotomy was performed in the left hemisphere, with a hand-held drill (burr tip 0.9 mm^2^); coordinates from bregma and midline: −2.8 from bregma, +4.2 from the midline. AAV injection into the EC was carried out through this craniotomy, at a depth of −2.8 to −2.7 mm from brain surface; the fiber-optic cannula positioning reached −2.6 mm from the surface. Coordinates were taken according to the Paxinos mouse brain atlas ^50^. The injection of adeno-associated virus (AAV) was performed in the target location using a precision pump (KD Scientific Inc., Harvard Bioscience) with a 10 μl NanoFil syringe with a 33-gauge beveled needle (NF33BV-2). The AAV used ^51^ (AAV5-CaMKIIa-hChR2(H134R)-mCherry (N_WT_ = 10, N_3xTgAD_ = 12), AAV5-CaMKIIa-mCherry (N_WT-mCherry_ = 9), titer 1-8×10^12^ vg/ml), were acquired from Vector Core at the University of North Carolina (USA). A total volume of 0.75 μl of the vector was injected in each mouse at a rate of 0.15 μl/min. To avoid backflow, the needle was kept in position for 10 minutes; after the needle extraction, a fiber optic cannula (diameter 200 μm, 0.39 NA, length according to the injection site, diameter 1.25 mm ceramic ferrule) was lowered to the targeted region (Laser 21 Pte Ltd, Singapore; Hangzhou Newdoon Technology Co. Ltd, China). The cannula was fixed in place with dental cement (Meliodent rapid repair, Kulzer). Buprenorphine was administered post-surgically to each animal. Animal recovery took place on a warm pad.

### Imaging: Animal preparation

Animal preparation followed a previously established protocol ^52^. Anesthesia was induced with 4% isoflurane; subsequently, the animals were endotracheally intubated, placed on an MRI-compatible cradle and artificially ventilated (90 breaths/minute; Kent Scientific Corporation, Torrington, Connecticut, USA). A bolus with a mixture of Pancuronium Bromide (muscle relaxant, Sigma-Aldrich Pte Ltd, Singapore) and Medetomidine (Dormitor, Elanco, Greenfield, Indiana, USA) was provided subcutaneously (0.05 mg/kg). A maintenance infusion (0.1 mg/kg/hr) was administered 5 minutes later, with isoflurane reduced to 0.5%.

Functional MRI was acquired 20 min following maintenance infusion onset to allow for the animal state to stabilize. Care was taken to maintain the temperature of the animals at 37 °C.

### Resting state functional-MRI

Data were acquired on an 11.75 T (Bruker BioSpin MRI, Ettlingen, Germany) equipped with a BGA-S gradient system, a 72 mm linear volume resonator coil for transmission. A 2 × 2 phased-array cryogenic surface receiver coil was adopted for the rs-fmri experiment (N = 29, respectively N_WT_ = 10; N_3xTgAD_ = 19; for a total of n = 52 runs) and a 10 mm single-loop surface coil for ofMRI experiments (N = 31, see below for details). Images were acquired using Paravision 6.0.1 software.

The parameters for the rs-fmri data acquisition are as follows: an anatomical reference scan was acquired using a spin-echo turboRARE sequence: field of view (FOV) = 17×9 mm^2^, FOV saturation slice masking non-brain regions, number of slices = 28, slice thickness = 0.35, slice gap = 0.05 mm, matrix dimension (MD) = 200×100, repetition time (TR) = 2750 ms, echo time (TE) = 30 ms, RARE factor = 8, number of averages = 2. Functional scans were acquired using a gradient-echo echo-planar imaging (EPI) sequence with the same geometry as the anatomical: MD = 90×60, TR = 1000 ms, TE = 15 ms, flip angle = 50°, volumes = 600, bandwidth = 250 kHz.

### Optogenetics functional-MRI

Parameters for the ofMRI data acquisition were adapted to the lower sensitivity of the room temperature receiver coil. The anatomical reference scan was acquired using FOV = 20 × 10 mm^2^, number of slices = 34, slice thickness = 0.35, slice gap = 0 mm, MD = 200 × 100, TR = 2000 ms, TE = 22.5 ms, RARE factor = 8, number of averages = 2. Functional scans were acquired using FOV = 17×9 mm^2^, FOV saturation slice masking non-brain regions, number of slices = 21, slice thickness = 0.45, slice gap = 0.05 mm, MD = 60 x 30, TR = 1000 ms, TE = 11.7 ms, flip angle = 50°, volumes = 720, bandwidth = 119047 Hz. Field inhomogeneity was corrected using MAPSHIM protocol. Light stimulation was provided through a blue light laser (473 nm, LaserCentury, Shanghai Laser & Optic Century Co., Ltd; ∼12-15 mW output with continuous light at the tip of the fiber) controlled by in-house software (LabVIEW, National Instruments).

After an initial 50 s of rest as a baseline, 10 ms light pulses were applied at 5, 10 or 20 Hz for 10 s followed by a 50 s rest period, in a 10-block design fashion. An additional 60 s of rest were recorded after the last block of stimulation. The experimental groups (3xTgAD and WT mice with ChR2-mCherry) and the negative control group (wild-type mice with mCherry alone) underwent the same imaging protocol, i.e., one resting-state scan, followed by 5 Hz, 10 Hz and 20 Hz evoked fMRI scans in randomized order. The negative control group was imaged with the same imaging protocol as the experimental groups to exclude potential heating and/or vascular photoreactivity artifacts ^53,54^. This resulted in a ofMRI dataset with N = 31 mice (N_WT-mCherry_ = 9, N_WT-ChR2_ = 10, N_3xTgAD-ChR2_ = 12), for a total of n = 147 runs. Additionally, in order to exclude abnormal behavior induced by the photostimulation protocol, all animals underwent the three stimulation sessions (5 Hz, 10 Hz, and 20 Hz) again while awake and freely walking in a behavior-chamber.

### Imaging: fMRI analysis

Images were processed using a protocol optimized for the mouse and corrected for spikes (*3dDespike, AFNI* ^55^, *motion (FSL mcflirt*, ^56^; and B1 field inhomogeneity (*FSL fast*). Automatic brain masking was carried out on the EPI using *FSL bet*, following smoothing with a 0.3 mm^2^ kernel (*FSL susan*) and a 0.01 Hz high-pass filter (*fslmaths*). Nuisance regression was performed using FIX. Separate classifiers were generated for rs-fmri and ofMRI based on 15 randomly-selected manually-classified runs from each study ^57^. The EPIs were registered to the Allen Institute for Brain Science (AIBS) reference template ccfv3 using SyN diffeomorphic image registration (*antsIntroduction.sh, ANTS* ^58^). The atlas resampled to 90 regions-of-interest, nomenclature, and abbreviations for the brain regions are in accordance with https://atlas.brain-map.org/.

### Whole-cell patch recording of acute brain slices

Mouse brains were rapidly removed after decapitation and placed in high sucrose ice-cold oxygenated artificial cerebrospinal fluid (ACSF) containing the following (in mM): 230 sucrose, 2.5 KCl, 10 MgSO_4_, 0.5 CaCl_2_, 26 NaHCO_3_, 11 glucose, 1 kynurenic acid, pH 7.3, 95% O_2_ and 5% CO_2_. Coronal brain slices were cut at a thickness of 250 mm using a vibratome (VT1200S; Leica Biosystems) and immediately transferred to an incubation chamber filled with ACSF containing the following (in mM): 119 NaCl, 2.5 KCl, 1.3 MgCl_2_, 2.5 CaCl_2_, 1.2 NaH_2_PO_4_, 26 NaHCO_3_, and 11 glucose, pH 7.3, equilibrated with 95% O_2_ and 5% CO_2_. Slices were allowed to recover at 32° C for 30 minutes and then maintained at room temperature. Experiments were performed at room temperature. Whole-cell patch-clamp recordings were performed on EC pyramidal cells expressing ChR2-mCherry and were visualized using a CCD camera and monitor. Pipettes used for recording were pulled from thin-walled borosilicate glass capillary tubes (length 75 mm, outer Ø 1.5 mm, inner Ø 1.1 mm, WPI) using a DMZ Ziets-Puller (Zeitz). Patch pipettes (2–4 MW) were filled with internal solution containing (in mM): 105 K-gluconate, 30 KCl, 4 MgCl_2_, 10 HEPES, 0.3 EGTA, 4 Na-ATP, 0.3 Na-GTP, and 10 Na_2_-phosphocreatine (pH 7.3 with KOH; 295 mOsm), for both voltage- and current-clamp recordings. Photostimulation (460 nm) was delivered by an LED illumination system (pE-4000). Several trains of a square pulse of 20 ms duration with 5, 10, and 20 Hz, were delivered respectively under current-clamp mode (I = 0) to examine whether the neurons were able to follow high-frequency photostimulation. After different frequencies of photostimulation were completed, neurons were shifted to voltage-clamp mode (at −60 mV), and a prolonged square pulse of 500 ms duration was delivered, to further confirm whether ChR2-induced current could be seen in the recorded neurons. The access resistance, membrane resistance, and membrane capacitance were consistently monitored during the experiment to ensure the stability and the health of the cell.

### Hidden Markov model of whole-brain BOLD fMRI fluctuations in response to optogenetic stimulation

BOLD fMRI time series were modelled using hidden Markov Models (HMMs) as a finite set of transient, repeating states. Each state is represented by a multivariate Gaussian distribution, which is described by the mean and covariance (respectively, BOLD amplitude and functional connectivity) ^48^. BOLD ofMRI time series were extracted from the AIBS atlas resampled to 90 regions-of-interest (https://atlas.brain-map.org/). HMMs were fit on a reduced space of principal components accounting for 50% of the variance in these time series.

For ofMRI data, we applied one single HMM on all concatenated subjects across experimental groups: healthy control group (N_WT-mCherry_ = 9), healthy active group (N_WT-ChR2_ = 10), and AD-like pathology active group (N_3xTgAD-ChR2_ = 12), thus a total of N = 31 mice, n = 153 runs. We applied the HMM with different states number (10, 12, 14, 16, and 18) and, for each number, we repeated the fitting 3 times, thus resulting in 3 HMMs per state number. We then identified the best HMM configuration for this ofMRI dataset as the number of states that provided the highest similarity between repetitions with lowest average free energy (for each state number, this was calculated as the ratio between model similarity and average free energy across repetitions). This allowed us to identify a remarkably robust temporal organization for 14 states (see **Fig. S5a**). Subsequent analyses were carried out on HMM with 14 states and lowest free energy value. We also show that HMMs with different states number were also capable of capturing brain dynamics induced by optogenetics (see **Fig. S7**).

Although the states were inferred at the group level, each animal has their own characteristic state time course representing the subject-specific probability of each HMM state being active at each instant (here, a fMRI volume). This is possible because we used variational Bayes in order to provide an analytical approximation of the model posterior distribution (see the original paper for more details ^1^). Statistical testing for inferences across groups or across stimulation frequencies, were carried out on these states’ time-series depicting time-resolved probability of activation. Details are provided below.

### Hidden Markov model of spontaneous whole-brain BOLD fMRI fluctuations during rest

For the rs-fmri dataset, BOLD time-series extraction and HMM fitting were performed as for the ofMRI dataset. As the primary aim of this work was to study the effect of optogenetic stimulation on brain network dynamics, for the analysis of rs-fmri data we fit a separate HMM now fixing the number of states to 14 in order to have the same order of dimensionality as for ofMRI data. Across mice, we then studied the spontaneous hierarchical organization in HMM states fractional occupancies (see below for details). We then also fit HMMs with different state numbers (10, 12, 14, 16, and 18) in order to study the robustness of the spontaneous hierarchical organization. HMMs on rs-fmri data were fit on N = 29 mice (N_WT_ = 10; N_3xTgAD_ = 19; for a total of n = 53 runs).

### Inference testing on HMM activation probabilities

All inference testing was carried out using Permutation Analysis of Linear Models (PALM: https://fsl.fmrib.ox.ac.uk/fsl/fslwiki/PALM, ^59^). The null distribution was characterized with 1,000 permutations. Statistical significance was established based on family-wise-error (FWE)-corrected p-value. When testing inferences on HMM responses to optogenetic stimulation (i.e. group comparisons: WT-ChR2 vs WT-mCherry), one-dimensional threshold-free cluster enhancement (TFCE) was applied, thus FWE-corrected p-values were fully corrected across time. Also, in exploratory analyses, statistical significance was further corrected across all states tested, thus p-values are FWE-corrected across time and states. All statistical significance results are plotted as −log_10_ of FWE-corrected p-values. All statistical analyses were carried out in MATLAB 2018.

### HMM response time-locked to optogenetic stimulation (short-scale)

Each ofMRI run comprised 10 stimulation blocks. In order to create HMM responses time-locked to stimulation, for each run, we considered the HMM activation probability 10 seconds from beginning of stimulation, plus 10 seconds after the end of stimulation, for a total of 20 seconds (10 on, 10 off). We then calculated the median across the 10 blocks - as this is less affected by outliers and skewed data and is usually the preferred measure when the distribution is not symmetrical. Short-scale HMM responses were studied by contrasting groups: WT-mCherry vs WT-ChR2, or WT-ChR2 vs 3xTgAD. Significance was assessed through permutation testing as explained above in “*Inference testing on HMM time-series*”. The results from this analysis can be found in **Fig. 2c, 5c**.

### Long-scale variation in short-scale HMM responses to optogenetic stimulation

HMMs characterize time-series for each state across the whole ofMRI run: we can therefore assess how short-scale HMM responses vary across long-scales. In order to do this in an unbiased fashion, we used permutation-based statistical significance as a criteria to deem a HMM response to a single block significant or not. Long-scale HMM responses were studied by contrasting WT-mCherry group vs WT-ChR2 group.

The results from this analysis can be found in **Fig. 3b**.

### Frequency effect on time-locked HMM responses

Each subject underwent multiple ofMRI runs at different frequencies (5, 10, and 20 Hz). In order to study the effect of frequency on time-locked HMM responses, a repeated measure ANOVA was carried out (as implemented in FSL randomise: https://fsl.fmrib.ox.ac.uk/fsl/fslwiki/Randomise). Given the key role of theta rhythms in the EC-hippocampal circuit, our hypotheses predicted specific responses to each theta rhythm (5 and 10 Hz). We therefore specifically contrasted 5 Hz to 10 and 20 Hz, and 10 Hz to 5 and 20 Hz, and carried out an overall F-test for the effect of stimulation frequency as well as post-hoc T-tests. Each set of three runs with frequencies at 5, 10, and 20 Hz (also referred in the manuscript as *session*), was considered as an independent block, and block-aware permutation testing was carried out (combination of within-block and whole-block) ^60^. Frequency effects on short-scale HMM responses were studied separately for WT-ChR2 group and for 3xTgAD group. The results from this analysis can be found (for WT-ChR2 group) in **Fig. 4a, S10b, g, S14c, e**, and (for 3xTgAD group) in **Fig. S13b, S14b, S14d, f**.

### Interaction effect of group by frequency on HMM responses

In order to study group by frequency interactions (theta rhythms) on time-locked HMM responses, a mixed-design ANOVA was carried out using FSL PALM (^59^, design matrix can be seen in **Fig. S14a**).

Group (WT vs 3xTgAD) was a between-subjects factor (fixed), while frequency (5 Hz to 10 and 20 Hz, and 10 Hz to 5 and 20 Hz) was a within-subject factor (random). An overall F-test of the interaction effect with both stimulation contrasts was carried out as well as post-hoc T-tests (for 5 Hz to 10 and 20 Hz: WT > 3xTgAD and 3xTgAD > WT; and, for 10 Hz to 5 and 20 Hz: WT > 3xTgAD and 3xTgAD > WT). Each set of three runs with frequencies at 5, 10, and 20 Hz (also referred in the manuscript as *session*), was considered as an independent block, and block-aware permutation testing was carried out (combination of within-block and whole-block) ^60^. The results from this analysis can be found in **Fig. 5e, f**.

We also tested a group by beta-frequency interaction on state 4 activation probability. Given we hypothesised greater activation probability for 3xTgAD compared to WT when driving stimulation at 20 Hz compared to 5 and 10 Hz, only one single T-test for the relevant interaction was tested. The results from this analysis can be found in **Fig. S13a**.

### General linear model of whole-brain BOLD fMRI fluctuations in response to optogenetic stimulation

Brain hemodynamic fluctuations during ofMRI were also examined using a traditional general linear model (GLM) framework (*FSL fsl_glm*). The stimulation paradigm and its first derivative were convolved using the default gamma function and used as regressors in the analysis, with motion parameters as covariates.

Threshold free cluster enhanced (TFCE) FWE-corrected p-values were deemed significant when below 0.05 after permutation testing. This analysis was carried out on WT-ChR2 group only. The results from this analysis can be found in **Fig. S8**.

### Multi-scale community detection

In resting-state fMRI BOLD data, and across multiple HMMs with varying numbers of states, we sought to identify clusters of HMM states sharing similar variance in FO in spontaneous brain network dynamics across mice. Across mice, we concatenated FO for HMMs with 10, 12, 14, 16, and 18 states, thus resulting in a meta-correlation matrix of FO similarity across mice. Then, a Louvain method for community detection was applied on this meta-correlation matrix. We performed a multi-scale community detection across a range of the structural resolution parameter γ ^61^. By maximizing the modularity quality function ^62^ using a Louvain-like ^63^ locally greedy algorithm (with 100 optimisations) for multiple values of γ, we quantitatively estimate consensus between partitions, calculating partitions similarity as z-score of the Rand coefficient ^64^. This analysis was carried out on the resting-state fMRI dataset only. The results from this analysis can be found in **Fig. 3b**.

### Directed acyclic graph

We modelled the relationship between optogenetic stimulation on/off-set and HMM states time-series (5 variables) as a directed acyclic graph (DAG). We characterized the optimal Bayesian network explaining these relationships using Greedy Equivalence Search (www.phil.cmu.edu/projects/tetrad/), which allowed us to establish statistical conditional dependencies (edges) between these variables—primarily to estimate direct causal connections between variables. The Linear, Non-Gaussian, Acyclic causal Models (LiNGAM) ^65^ approach was then used to determine further the direction of the edges, i.e. the causality between the variables. Importantly, optogenetic stimulation on/off-set and HMM states time-series were treated equally with no further supervised labelling, thus the model was agnostic to what source generated the activity. Model comparison across all possible permutations was carried out using a Bayesian information criteria (BIC). This analysis was carried out on WT-ChR2 group only. The results from this analysis can be found in **Fig. 3c, 4d**.

### Association with EC monosynaptic projection

The pattern of EC monosynaptic tracing projection was derived from the publicly available data of the Allen Brain Institute ^66^. Magnitude values were normalized via rank-based inverse normal transformation.

Statistical association between EC projection pattern and spatial activity pattern of HMM states of interest was carried out using PALM ^59^. The null distribution was characterized with 1,000 permutations. Statistical significance was established based on FWE-corrected p-value. The results from this analysis can be found in **Fig. S7**.

### Structural-to-functional mapping

Mapping of anatomical projections derived from connectivity tracing on functional activation patterns was carried out using cross-validated regularized Ridge regression implemented in FSLNets (https://fsl.fmrib.ox.ac.uk/fsl/fslwiki/FSLNets). A visualization of the method can be found in **Fig. S11**. For all the regions considered in the functional brain analysis, the monosynaptic projection patterns were derived from the publicly available data of the Allen Brain Institute ^66^ into a brain region by brain region matrix. Predictor variables, with columns being different brain regions and, for each column, each row value being the magnitude of tracing projection to a target region, were normalized via rank-based inverse normal transformation and used to predict HMM state amplitude levels across brain regions (response variable). We carried out leave-one region-out cross-validation. The lambda value (Lagrange multiplier) was estimated through an inner loop. Statistical significance was assessed with 1,000 permutations. Estimated regression coefficients were then extracted and averaged across folds. These values were then dot-multiplied by the whole-brain directed connectivity matrix in order to characterize a directed connectivity matrix underlying a specific state of activity. To aid interpretation, only positive connections were considered and visualized as a circle graph. The results from this analysis can be found in **Fig. 4c, S10c-e, h-l**.

## Supplementary figures

**Figure S1.**
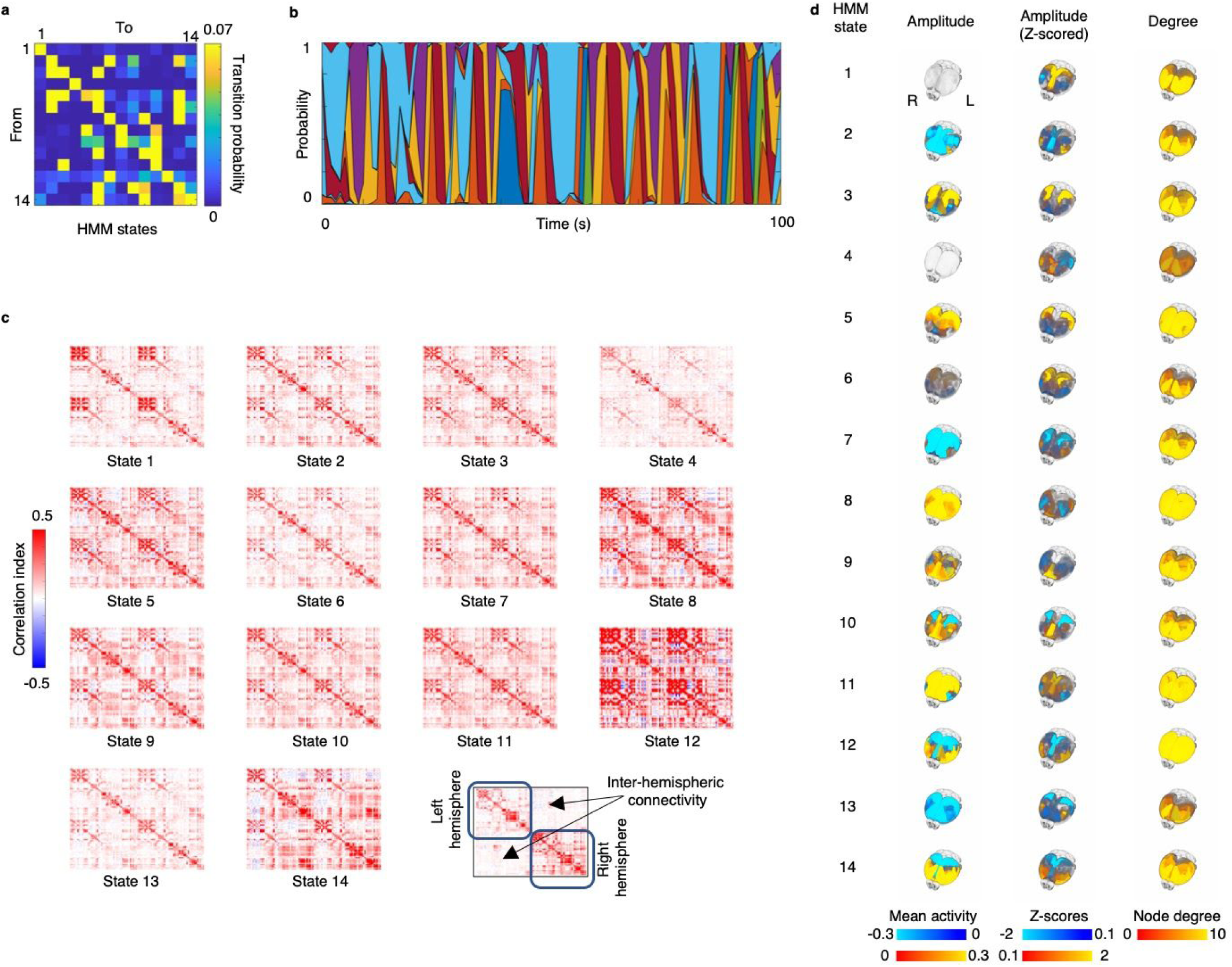
Spontaneous brain states of whole brain hemodynamic activity. HMM were used to study spontaneous brain network dynamics in the rs-fmri dataset (N = 29 mice (WT: N = 10; 3xTgAD, N = 19), n = 53 runs acquired over two sessions). Mice brain activity could be characterized by fast transitions between 14 different states of activity in a stochastic non-random pattern. **a**) Directed matrix of state-to-state transition probability, where directed transitions *from* a state *to* the others is represented by a matrix row. **b**) HMM states activation probability for some 100 seconds in a representing animal; each color represents a state. Brain network states are characterized as dynamics in both BOLD amplitude and functional connectivity. **c**) HMM functional connectivity pattern for each of the 14 HMM states. **d**) Showing HMM state maps as 3D brain images (for visualization and interpretation purpose, functional connectivity **e**) was transformed into node degree).

**Figure S2.**
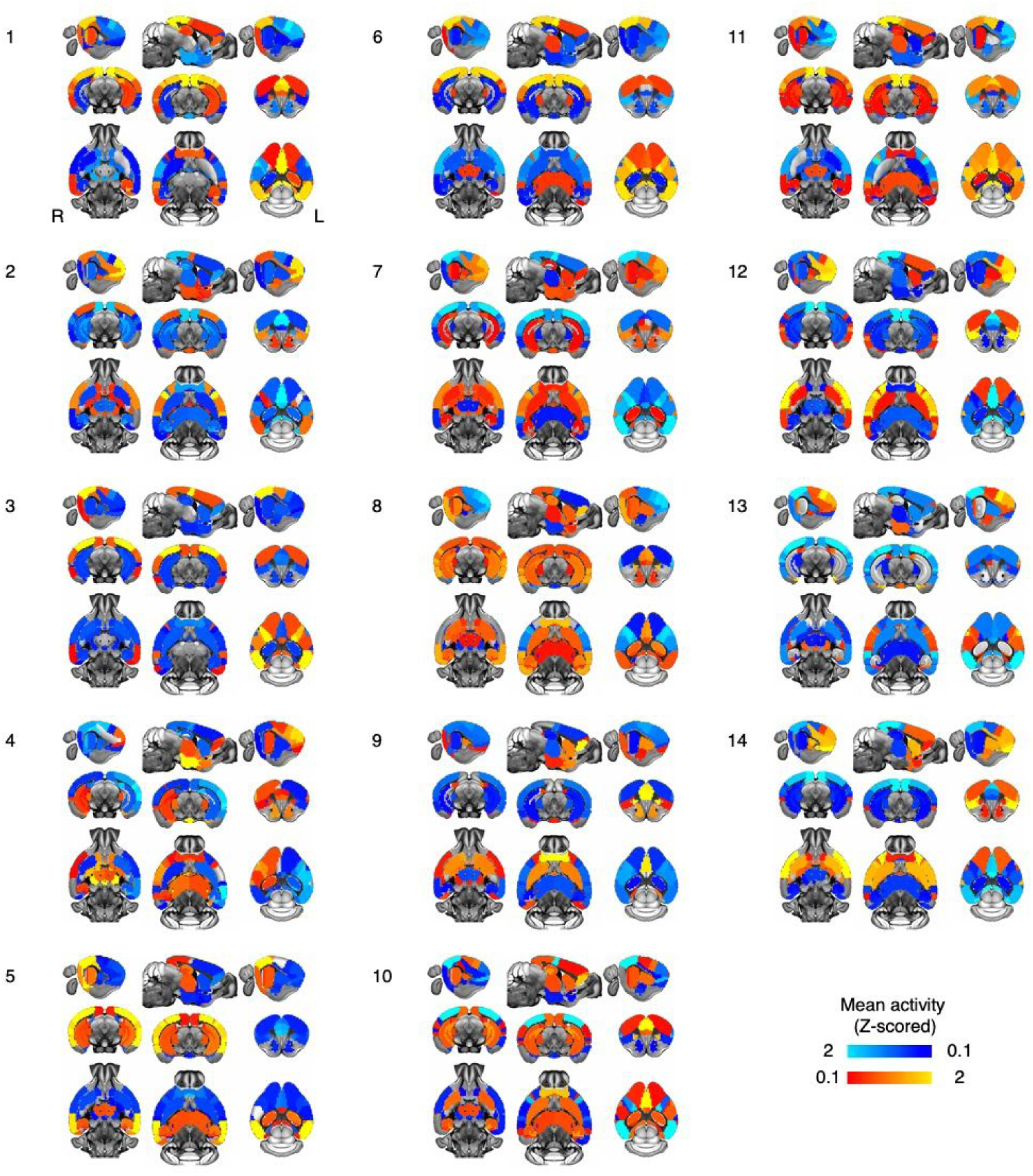
Maps of spontaneous HMM states of brain activity. 2D visualization of the 14 HMM states mean activity maps estimated in the rs-fmri dataset.

**Figure S3.**
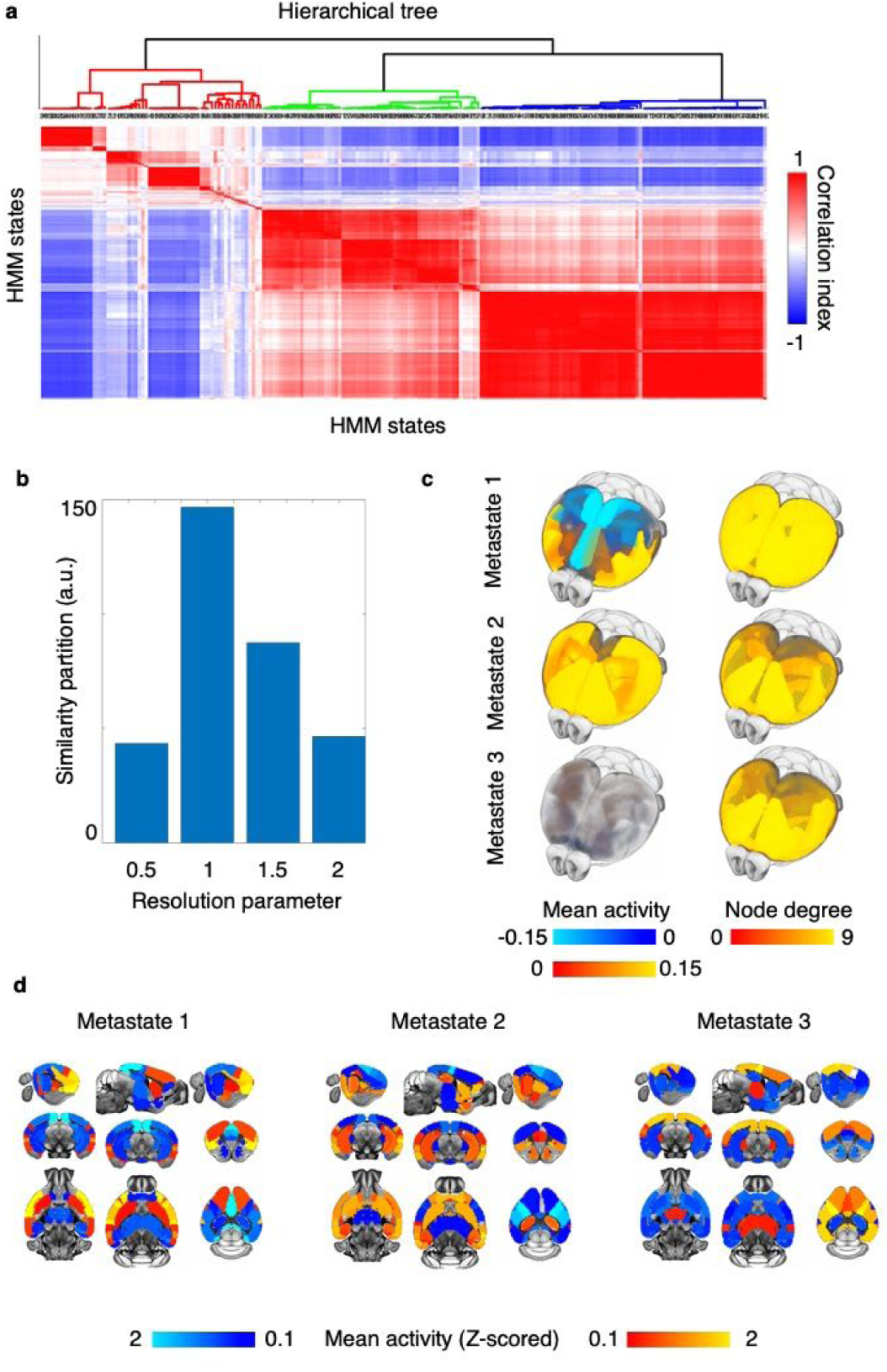
Spontaneous hemodynamic activity in the mouse brain is spatially and temporally organized. In the rs-fmri dataset (N = 29 mice (WT: N = 10; 3xTgAD, N = 19), n = 53 runs acquired over two sessions), HMMs with varying states numbers (10, 12, 14, 16, and 18) were assessed. For each configuration, states fractional occupancy (FO) was estimated : the amount of time spent in a state. **a**) Showing FO correlation matrix across mice for multiple HMMs with different numbers of states (10, 12, 14, 16, 18; concatenated together for a total of 70 states). In a data-driven fashion, multi-scale community detection was used in order to estimate the most robust number of communities of states or metastates across 100 repetitions. **b**) Showing mean partition similarity across a range of values for the resolution parameter. Maximum mean partition similarity was achieved for γ = 1. **c**) 3D representation of mean amplitude and node degree for metastates. d) 2D visualization of metastates mean amplitude.

**Figure S4.**
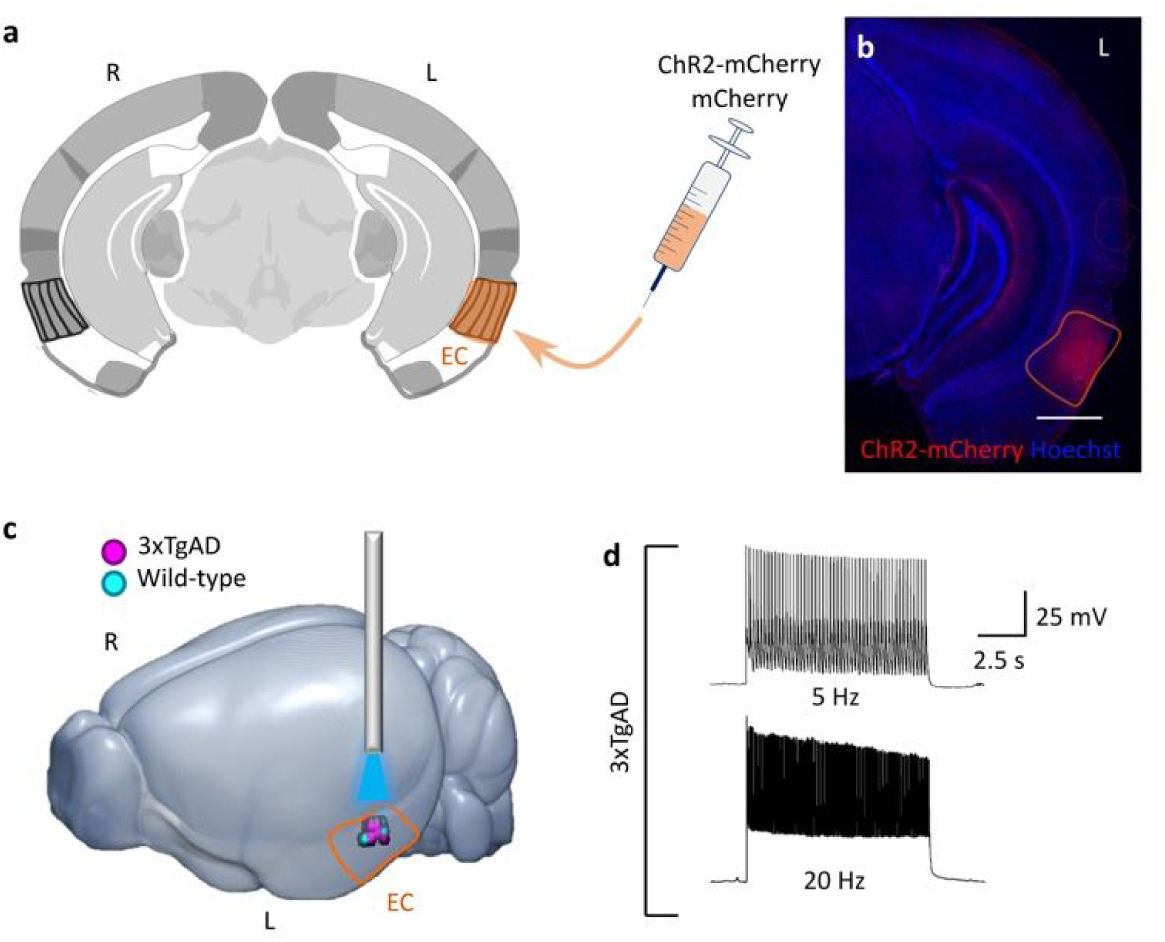
Optogenetic stimulation of excitatory neurons in the EC. **a)** Schematic illustrating CaMKII-ChR2-mCherry and CaMKII-mCherry transfection in the left EC. Coordinates relative to bregma and midline: −2.8 mm from bregma and +4.2 mm from the midline; depth of the injection *in loco:* −2.8/-2.6 mm. **b)** Histological validation confirms ChR2-mCherry expression (red) in the EC, with counter-stain with DAPI (blue) for cell nucleus. **c)** 3D rendering of the optrode stimulation targets clustered within the EC in both 3xTgAD (pink dots) and wild-type control mice (cyan dots). **d)** Transfected EC neurons, here shown in 3xTgAD, faithfully respond to 5 Hz and 20 Hz photostimulation pulses in ex-vivo slice recordings.

**Figure S5.**
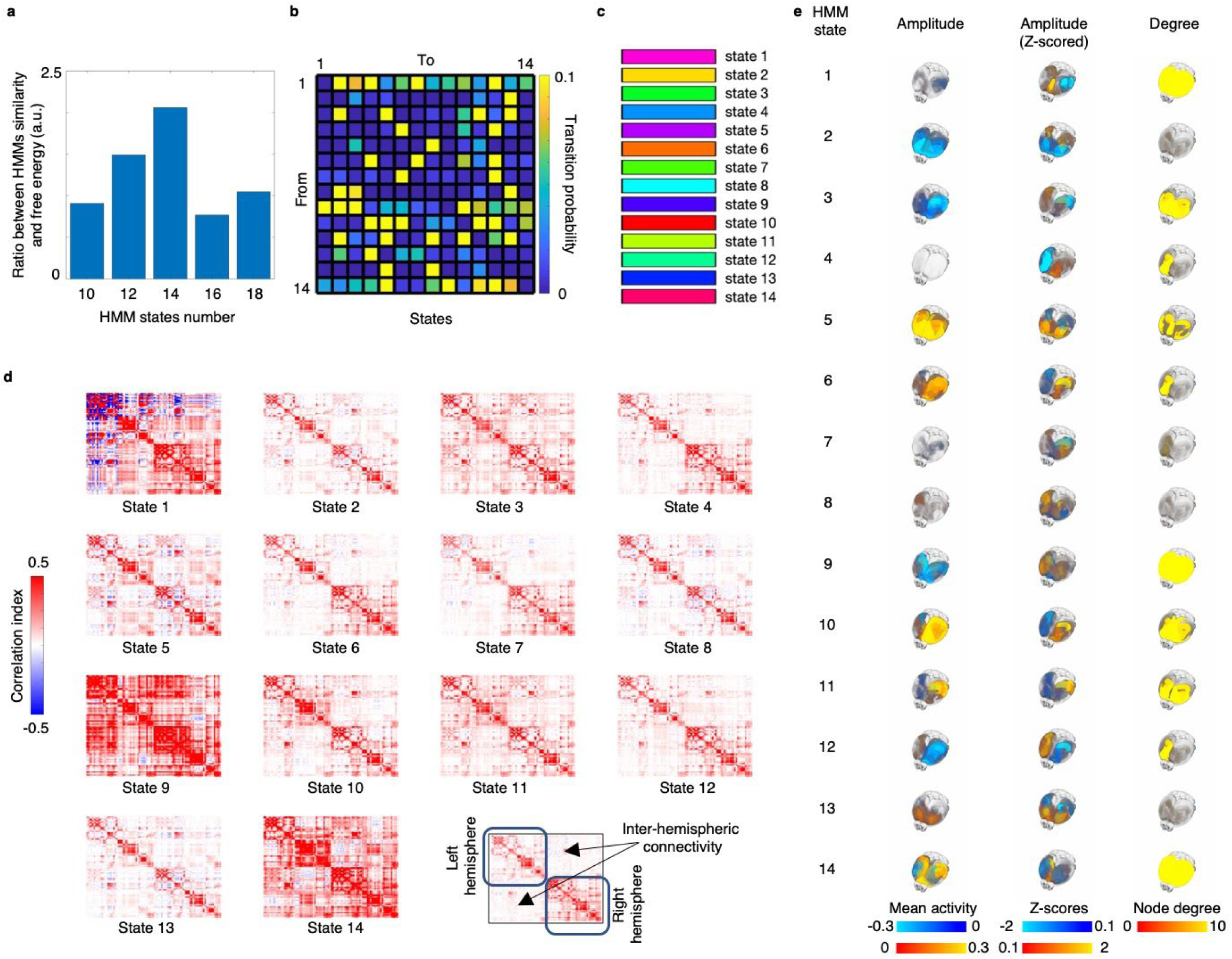
Brain states of whole brain hemodynamic activity in response to optogenetic stimulation. Hidden Markov models (HMMs) were used in order to characterize fast variations in whole brain BOLD fMRI fluctuations. Multiple HMM were fitted with different states numbers (10, 12, 14, 16, and 18 states) across all subjects and runs. Each separate HMM was fit on N = 31 mice, n = 153 runs (N_WT-mCherry_ = 9 mice, n = 27 runs acquired in 1 session; N_WT-ChR2_ = 10 mice; n = 54 runs acquired into 2 sessions; N_3xTgAD-ChR2_ = 12 mice; n = 72 runs acquired in two session; total: N = 31 mice, n = 153 runs). Each HMM state number configuration was assessed 3 times in order to calculate model similarity. The best HMM state number was identified based on the ratio between model similarity and average free energy. Results are shown in **a**): higher similarity would push the bar higher, as well as lower free energy. A state number of 14 led to models with greater similarity and lower free energy. For HMM with 14 states and lowest free energy: **b**) Showing matrix of transition probability between states. **c**) Showing color coding for HMM states used across all main figures picturing results from ofMRI. **d**) HMM functional connectivity pattern for each of the 14 HMM states. **e**) Showing HMM state maps as 3D brain images (for visualization and interpretation purpose, functional connectivity d) was transformed into node degree).

**Figure S6.**
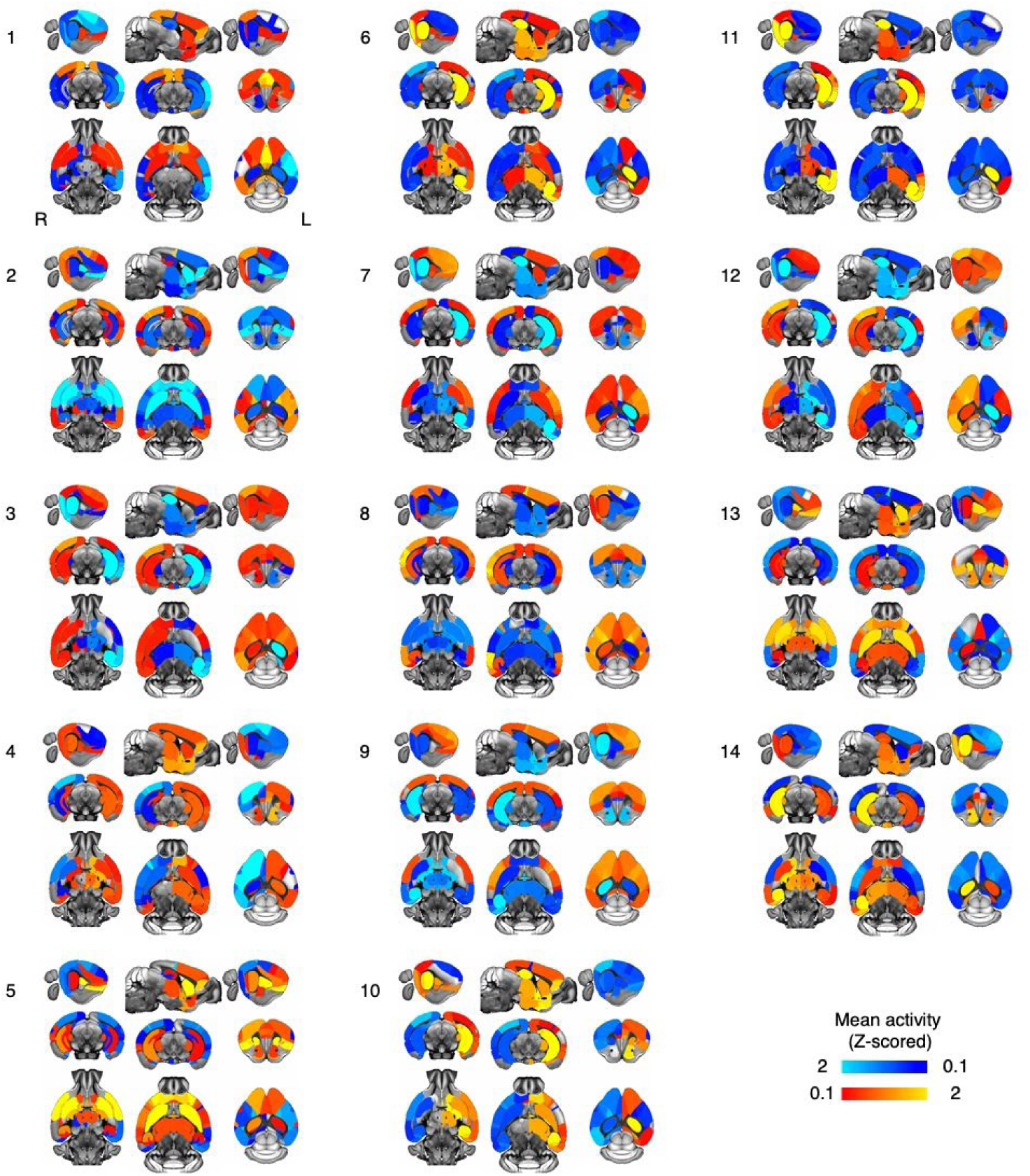
Maps of optogenetically-induced HMM states of brain activity. 2D visualization of the 14 HMM states mean activity maps estimated in the ofMRI dataset.

**Figure S7.**
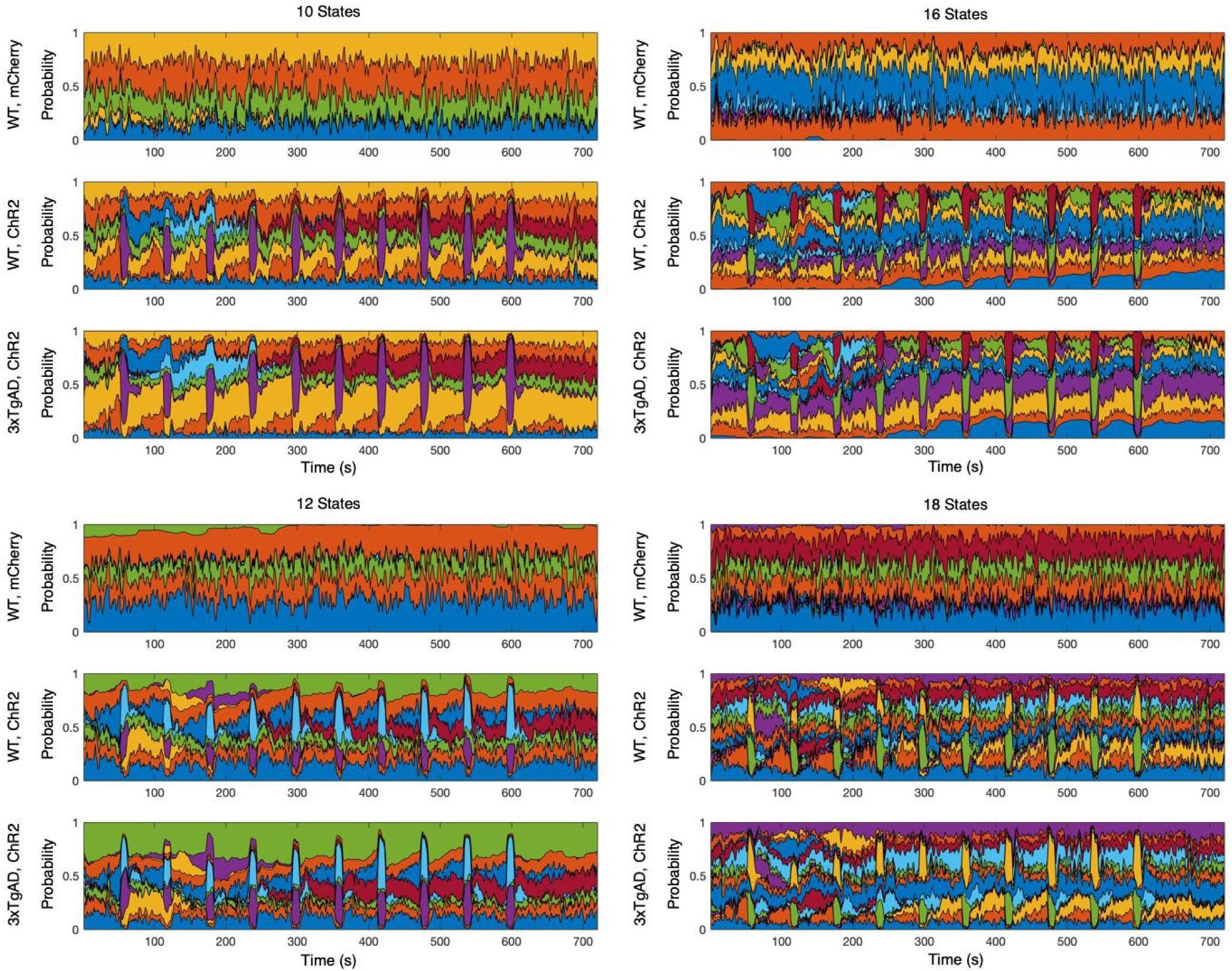
HMMs with different states number all show robust responses to optogenetic stimulation blocks. HMMs were fit with different state numbers. Although the HMM with 14 states showed greater similarity over model fitting repetitions and lower free energy, thus representing the best model, here HMM sensitivity to depicting BOLD changes to optogenetic stimulation across different state numbers is shown. Across all models, the HMMs were able to depict “bursts” in BOLD activity in response to stimulation, in both wild healthy mice and transgenic mice (both ChR2). Crucially these fast changes were not present in controls (mCherry). WT = wild-type. 3xTgAD = transgenic model of Alzheimer’s disease. Each separate HMM was fit on N = 31 mice, n = 153 runs (N_WT-mCherry_ = 9 mice, n = 27 runs acquired in 1 session; N_WT-ChR2_ = 10 mice; n = 54 runs acquired into 2 sessions; N_3xTgAD-ChR2_ = 12 mice; n = 72 runs acquired in 2 session; total: N = 31 mice, n = 153 runs).

**Figure S8.**
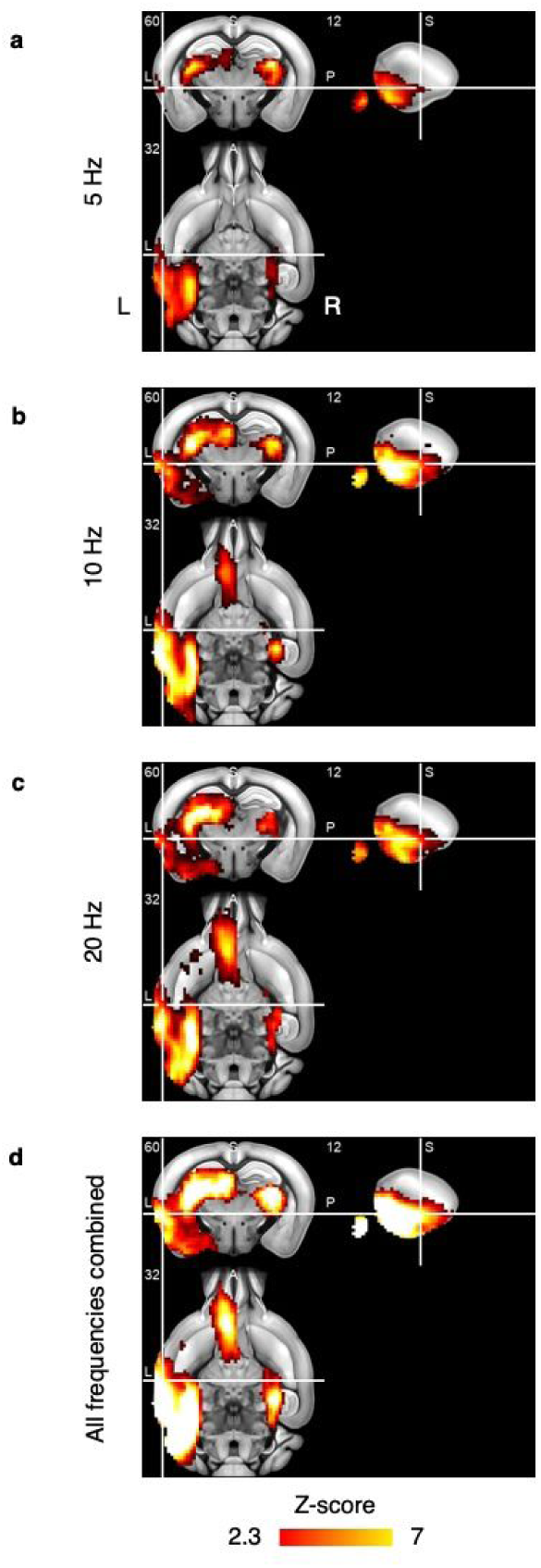
GLM modeling of BOLD fMRI response to optogenetic stimulation. Results from analyzing only the WT-ChR2 group (N_WT-ChR2_ = 10 mice; n = 54 runs; each animal underwent 2 sessions of 3 runs, one run per stimulation frequency (5, 10, 20 Hz)). Here, stimulation frequencies were collapsed together as well as explicitly modelled. Brain hemodynamic fluctuations during ofMRI were also examined using a traditional general linear model (GLM) framework. Here showing 2D brain maps depicting activation as Z-statistic level, thresholded at FWE-corrected p-values < 0.05. Showing, respectively, brain activation maps for stimulating EC at **a**) 5 Hz, **b**) 10 Hz, **c**) 20 Hz, and **d**) for all frequencies combined.

**Figure S9.**
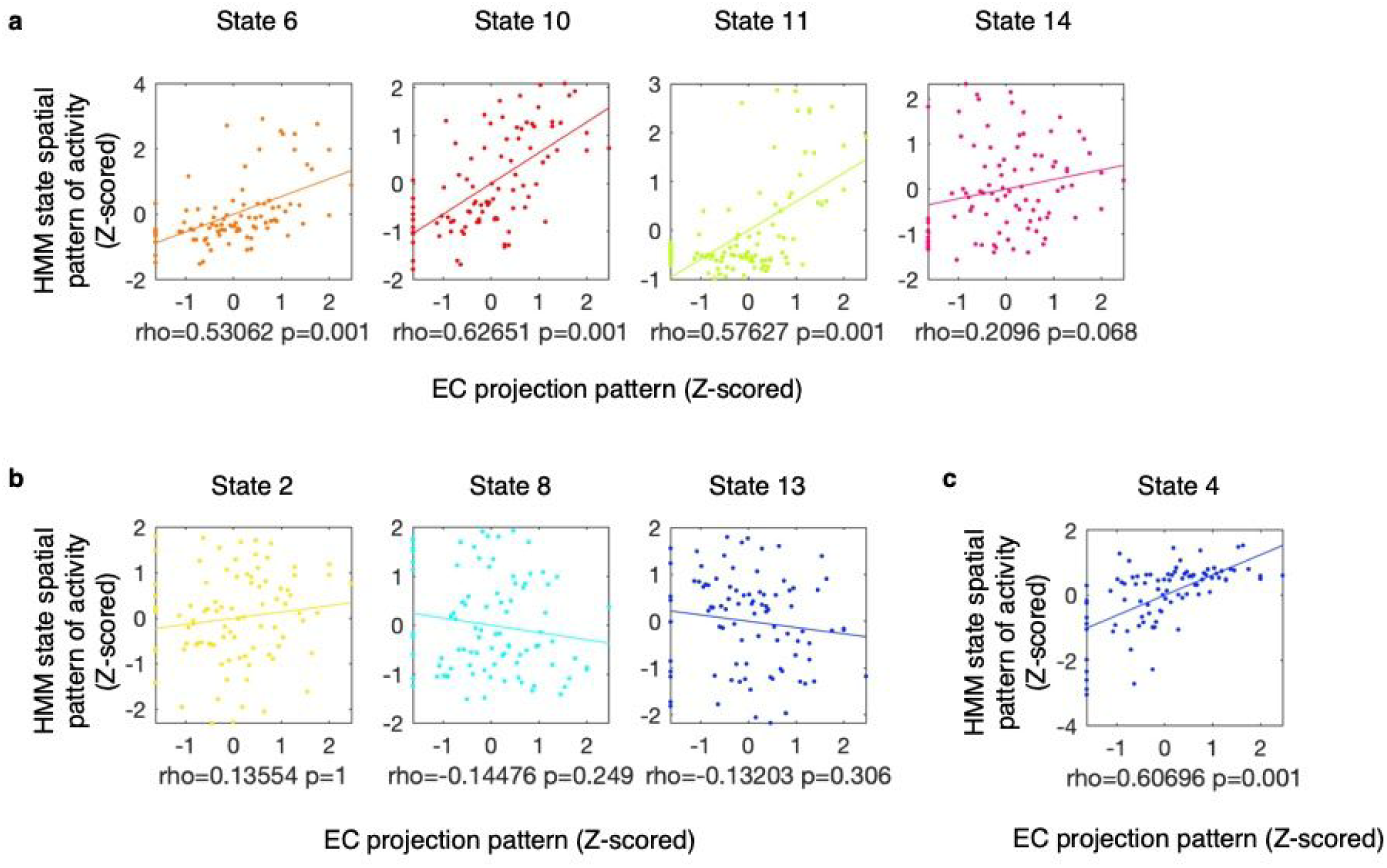
Association between EC monosynaptic projection and spatial pattern of HMM states mean activity. Showing scatter plots of association between EC monosynaptic projection pattern and spatial pattern of HMM states mean activity. EC projection pattern was normalized via rank-based inverse normal transformation and the association with the spatial pattern of HMM states mean activity was tested through FSL PALM. Rho-value indicates Pearson’s correlation score; p-value indicates FWE-corrected significance. **a**) States found to be significantly *more* active in WT-ChR2 group compared to WT-mCherry group. These states were tested for a positive association. **b**) States found to be significantly *less* active in WT-ChR2 group compared to WT-mCherry group. These states were tested for a negative association. **c**) Single state found to be significantly more active in the 3xTgAD group compared to the WT-ChR2 group. For this state a two-tailed test was carried out.

**Figure S10.**
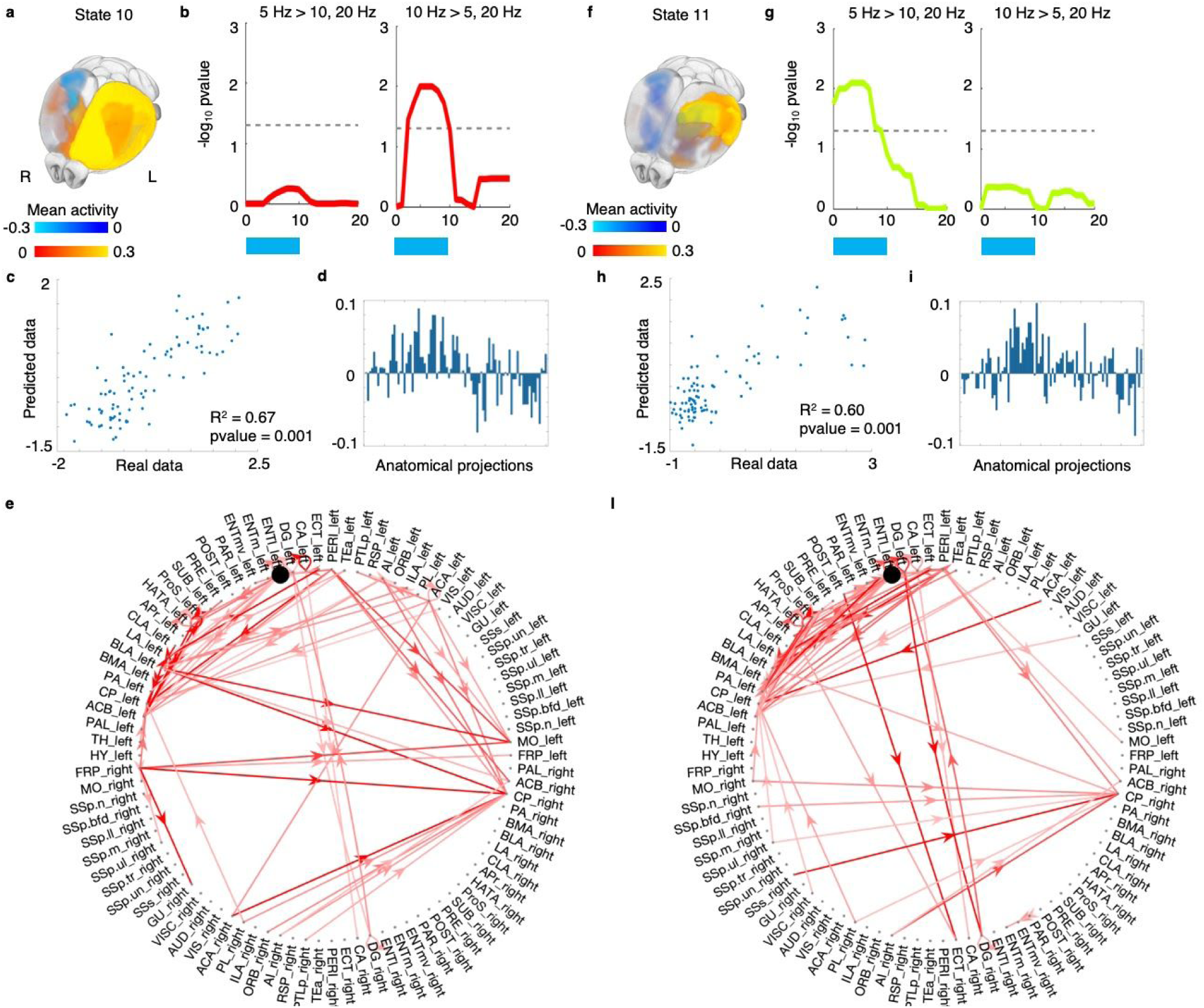
Different anatomical circuits predict activity patterns in distinct HMM states. Results from analyzing only the WT-ChR2 group (N_WT-ChR2_ = 10 mice; n = 54 runs; each animal underwent 2 sessions of 3 runs, one run per stimulation frequency (5, 10, 20 Hz)). Stimulation frequencies were explicitly modelled in the statistical tests. **a**) HMM mean activity map for state 10. **b**) showing results of repeated measures ANOVA post-hoc T-tests (as −log_10_ FWE-corrected p-values) comparing activation probability at 5 or 10 Hz with response at 20 Hz. Cross-validated Ridge regression (See **Fig. S10**) was then used together with publicly available tracing data (see Supplementary methods) to predict HMM mean activity map of state 10. **c**) Scatter plot of real vs predicted values for state 10 mean activity values. Average regression coefficients across folds were then extracted **d**) and plotted its positive values as a directed network (circular graph) **e**). The same approach was carried out for HMM state 11. Panels **f**), **g**), **h**), **i**), **l**) respectively mirror panels **a**), **b**), **c**), **d**), **e**) for *state 11*.

**Figure S11.**
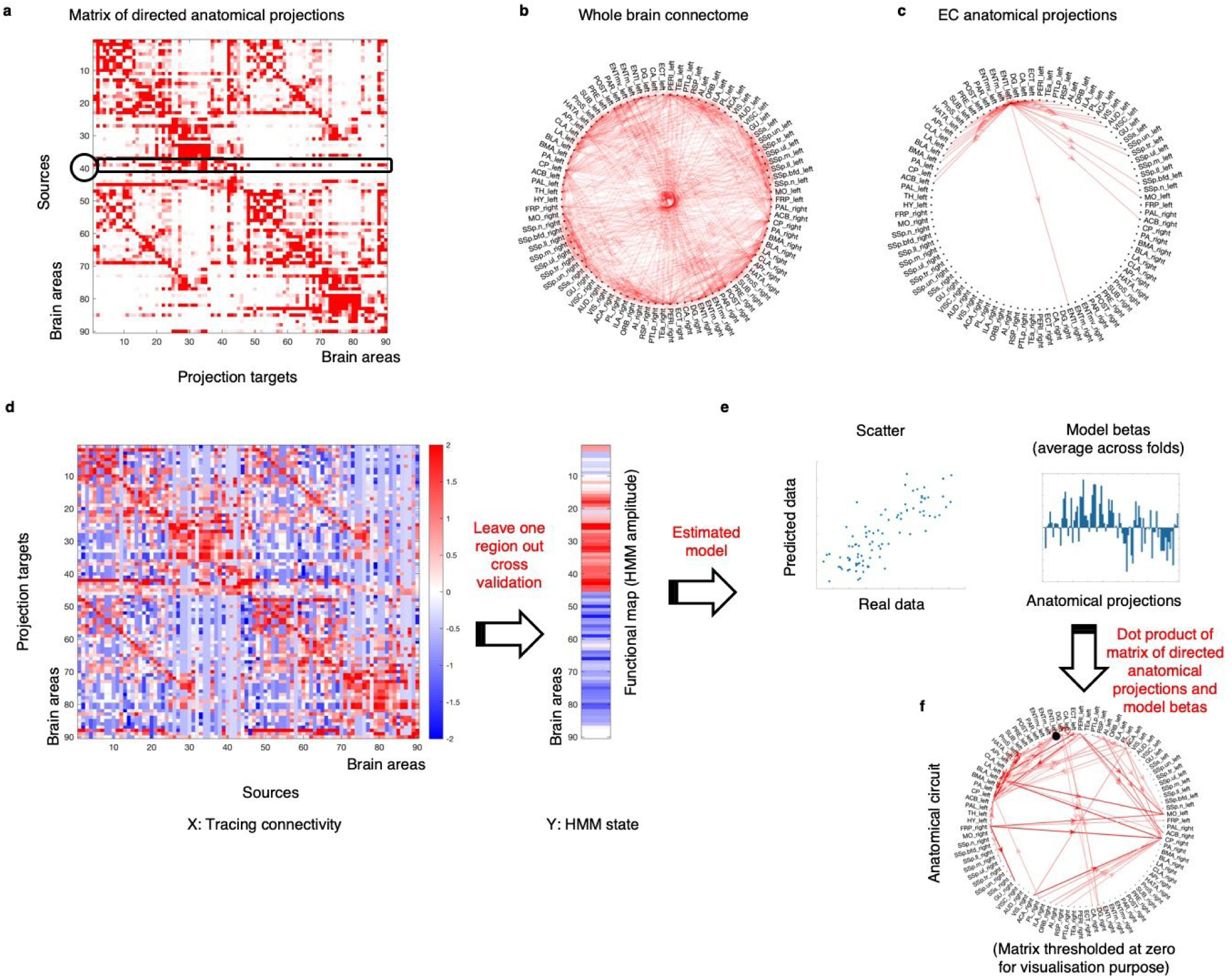
Structural to functional mapping. **a**) Monosynaptic tracing projection patterns were derived from the publicly available data of the Allen Brain Institute in the form of a brain region x brain region matrix. Each row represents the outgoing projection magnitude for one single brain region (source) to all other regions (target projections). **b**) Representation of A into a circular graph (thresholded only for visualization purpose). **c**) Outgoing connection from the left lateral EC. The matrix of directed anatomical projections (shown in A) was then transposed and normalized (see Supplementary methods) and used in a leave-one region-out cross-validation approach to predict functional maps of HMM activity amplitude. **d**) Showing a schematic carton of the data used in the regularized regression model. **e**) The estimated model, in terms of regression betas, were then averaged across cross-validation folds. Each beta represents the weight of one anatomical projection pattern in predicting function. These betas were then dot-multiplied by the directed anatomical projections matrix (shown in A) to provide a weighted anatomical circuit underlying the predicted HMM state. **f**) Showing the estimated anatomical circuit in the form of a circular graph.

**Figure S12.**
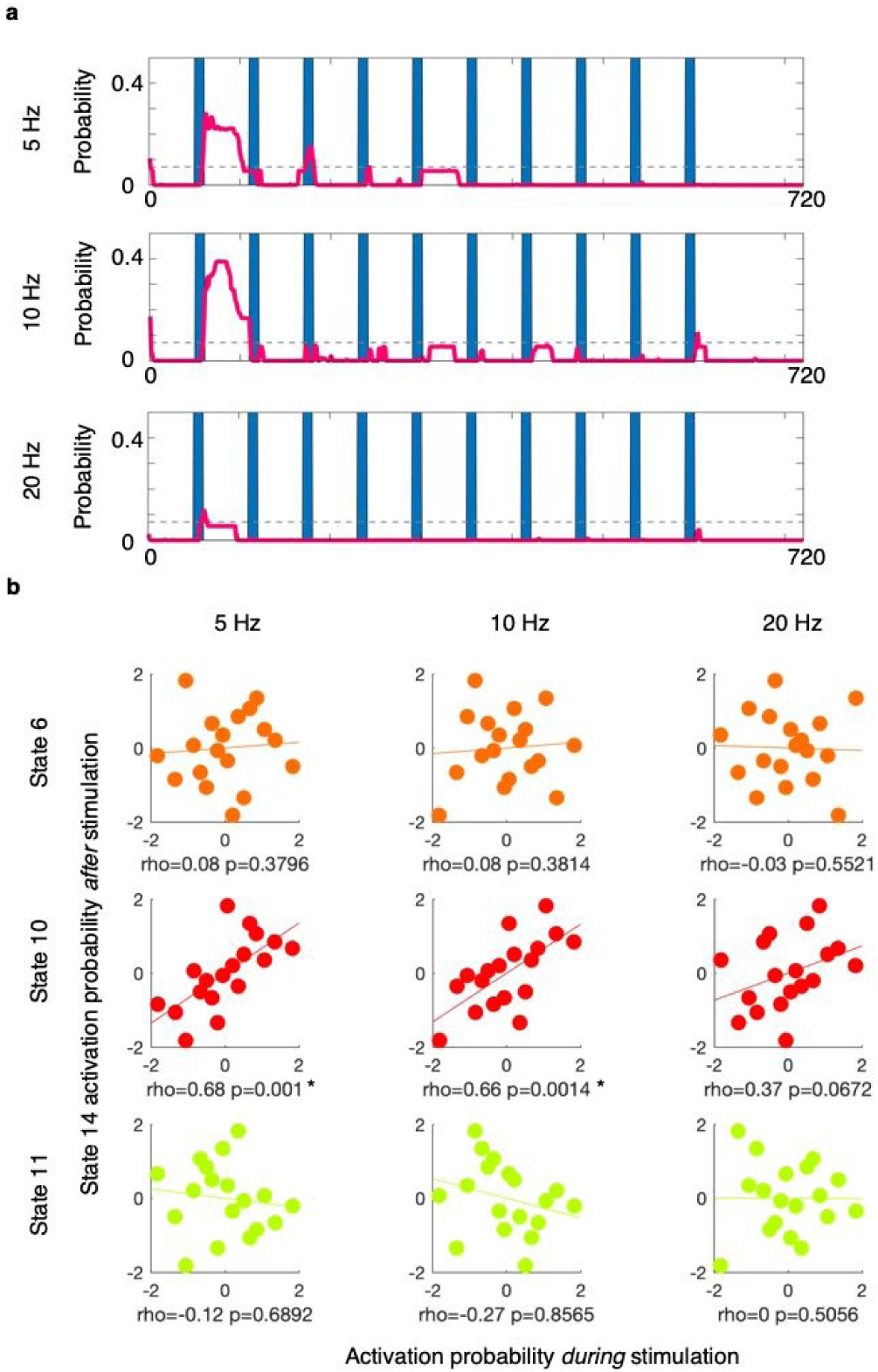
Downstream activity is caused by driving upstate brain network response with theta rhythms. Results from analyzing only the WT-ChR2 group (N_WT-ChR2_ = 10 mice; n = 54 runs; each animal underwent 2 sessions of 3 runs, one run per stimulation frequency (5, 10, 20 Hz)). Here, stimulation frequencies were explicitly modelled in the statistical tests. Optogenetic stimulation of EC also resulted in a downstream activity state (state 14). **a**) For each stimulation frequency, showing activation probability of HMM state 14 across stimulation blocks. As 1) theta oscillations are known to play a key role in the EC-hippocampal circuit, and 2) analyses shown in **Fig. 4** show that activation of state 14 is mediated by other states, here we asked whether, across mice, activity level in state 14 *after* stimulation is predicted by optogenetically-driven states responses *during* stimulation specifically within the theta rhythms. **b**) Scatter plot X axis shows state (6, 10, or 11) activation probability *during* stimulation; Y axis shows state 14 activation probability *after* stimulation. (Showing normalized values across subjects). Reported are Spearman’s correlation values (rho and p-value for a right tail test; Bonferroni corrected p-value = 0.0056; * significant after Bonferroni correction). Only state 10 activation level during stimulation with theta rhythms (5 and 10 Hz) showed to significantly correlate with state 14 activation level after stimulation.

**Figure S13.**
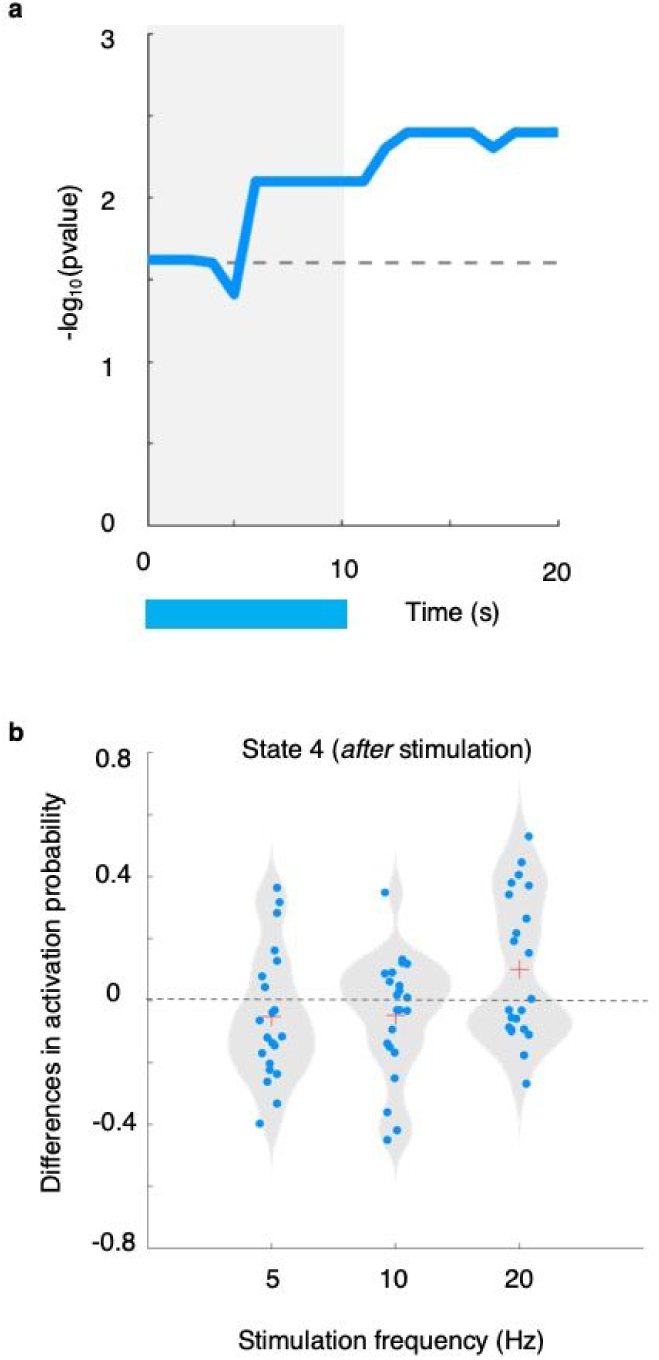
Positive effect of theta rhythms in AD mice outlast stimulation duration. Showing results of group by beta-frequency interaction between the WT-ChR2 group and the 3xTgAD-ChR2 group (WT-ChR2 group and 3xTgAD-ChR2 group (N_WT-ChR2_ = 10 mice; n = 54 run; N_3xTgAD-ChR2_ = 12 mice; n = 72 runs; for both groups, each animal underwent 2 sessions of 3 runs, one run per stimulation frequency (5, 10, 20 Hz)). **a**) Showing −log_10_ FWE-corrected p-values of T-test testing for greater activation probability in 3xTgAD compared to WT when inducing stimulation at 20 Hz compared to 5 and 10 Hz. These results indicate that in AD-like mice theta rhythms decreased state 4 activation probability significantly more than in WT mice, compared to beta rhythms. We then asked whether this effect outlasted stimulation duration, thus *after* stimulation ended. A repeated measure ANOVA in AD mice only was used to test the effect of theta rhythms (N_3xTgAD-ChR2_ = 12 mice; n = 72 runs; each animal underwent 2 sessions of 3 runs, one run per stimulation frequency (5, 10, 20 Hz)). **b**) Showing subject-specific difference in state 4 activation probability *after* stimulation. Each dot is a ofMRI run. Red cross indicates distribution mean. An overall effect (F-test) of frequency was found on state 4 activation probability after stimulation (F-test: p = 0.0100). Activation probability was found significantly reduced for both 5 and 10 Hz stimulation (post-hoc T-tests: respectively, 5 Hz stimulation, p = 0.0130; and 10 Hz stimulation, p = 0.0020; significance assessed with 1,000 permutations).

**Figure S14.**
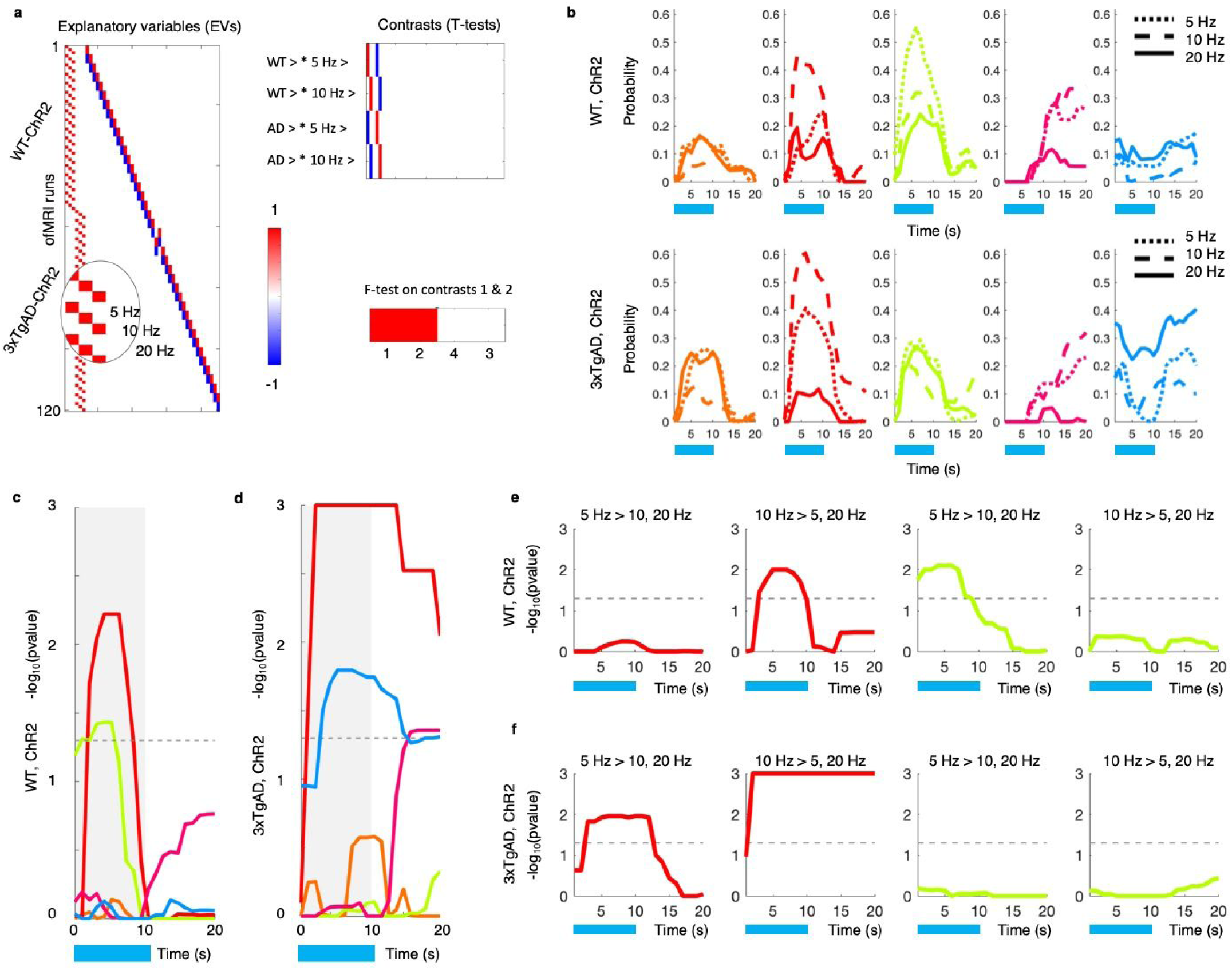
Post-hoc tests showing a group by frequency interaction in HMM response. Showing results of group by beta-frequency interaction between the WT-ChR2 group and the 3xTgAD-ChR2 group (WT-ChR2 group and 3xTgAD-ChR2 group (N_WT-ChR2_ = 10 mice; n = 54 run; N_3xTgAD-ChR2_ = 12 mice; n = 72 runs; for both groups, each animal underwent 2 sessions of 3 runs, one run per stimulation frequency (5, 10, 20 Hz)). **a**) Design matrix used to carry out a mixed-design ANOVA testing for a group *by* frequency interaction. **b**) Average HMM activation probability time-locked to optogenetic stimulation and divided by *group* (top row for the WT-ChR2 group, bottom for the 3xTgAD-ChR2 group) and *frequency* (dotted line: 5Hz; dashed line: 10Hz; solid line: 20 Hz). To confirm the results of the interaction analysis showed in **Fig. 5e, f**, two separate repeated measure ANOVA were carried out independently in the WT-ChR2 group (N_WT-ChR2_ = 10 mice; n = 54 run) and in the 3xTgAD-ChR2 group (N_3xTgAD-ChR2_ = 12 mice; n = 72 runs). **c**) Showing −log_10_ FWE-corrected p-values of F-test for effect of theta frequencies on in the WT-ChR2 group; results from post-hoc T-tests are shown in **e**). Plot c) is also shown in Fig. 4a. Here is reported again for direct comparison with the 3xTgAD-ChR2 group. **d**) Showing −log_10_ FWE-corrected p-values of F-test for effect of theta frequencies on in the 3xTgAD-ChR2 group; results from post-hoc T-tests are shown in **f**). Dotted line in **c-f** indicates statistical threshold.

## Supplementary tables

**Supplementary table 1.** Comparing correlations in predicting activity pattern of HMM state 10. Tracing projection data was used in a cross-validated fashion to predict the pattern of activity of state 10 and 11 and to estimate the respective regression betas representing the underlying anatomical circuits (respectively for state 10 and 11). Regression betas trained in state *11* prediction were then used to predict state 10 activity pattern. The table is showing the correlation values between the real pattern of state 10 activity and 1) predicted activity pattern based on state 10 prediction betas, and 2) predicted activity pattern based on state 11 prediction betas.

| Variable X | Variable Y | Pearson's rho |
| --- | --- | --- |
| Prediction betas state 10 to predict state 10 functional pattern | Activity pattern HMM state 10 | 0.82 |
| Prediction betas state 11 to predict state 10 functional pattern | Activity pattern HMM state 10 | 0.72 |
| Prediction betas state 10 to predict state 10 functional pattern | Prediction betas state 11 to predict state 10 functional pattern | 0.79 |

**Supplementary table 2.** Comparing correlations in predicting activity pattern of HMM state 11. Tracing projection data was used in a cross-validated fashion to predict the pattern of activity of state 10 and 11 and to estimate the respective regression betas representing the underlying anatomical circuits (respectively for state 10 and 11). Regression betas trained in state *10* prediction were then used to predict state 11 activity pattern. The table is showing the correlation values between the real pattern of state 11 activity and 1) predicted activity pattern based on state 11 prediction betas, and 2) predicted activity pattern based on state 10 prediction betas.

| Variable X | Variable Y | Pearson's rho |
| --- | --- | --- |
| Prediction betas state 11 to predict state 11 functional pattern | Activity pattern HMM state 11 | 0.77 |
| Prediction betas state 10 to predict state 11 functional pattern | Activity pattern HMM state 11 | 0.66 |
| Prediction betas state 11 to predict state 11 functional pattern | Prediction betas state 10 to predict state 11 functional pattern | 0.75 |

**Supplementary Table 3.**
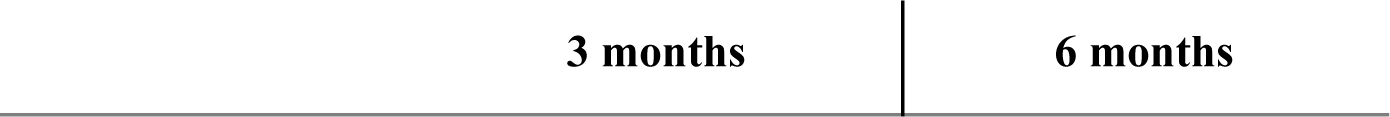
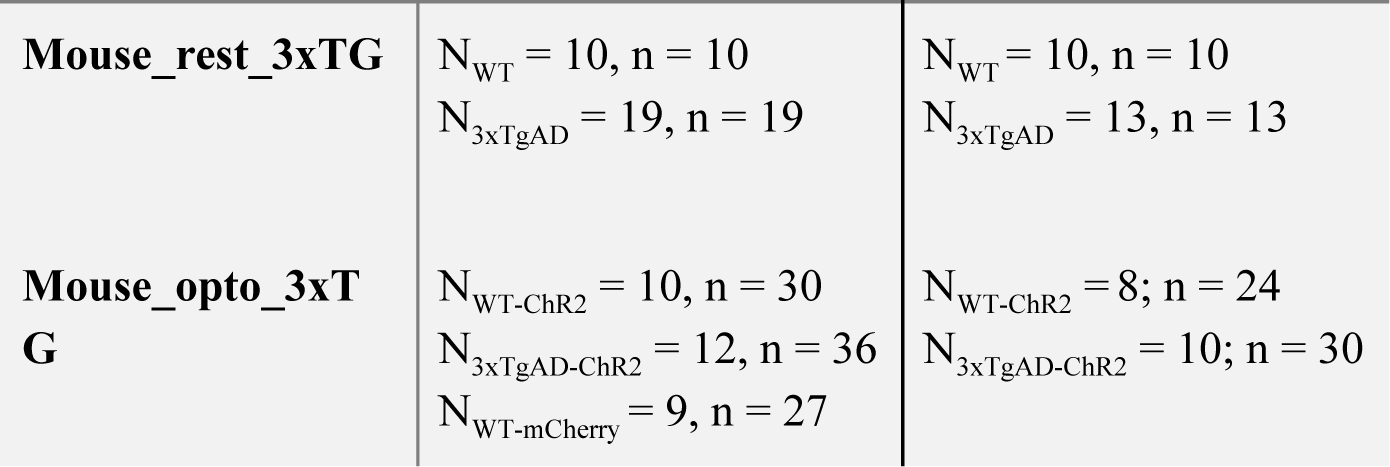
Animal breakdown per experiment type. Animal breakdown per experimental setting. rs-fmri was conducted on WT and 3xTgAD mice at 3 and 6 months of age, 1 run per mouse per time-point. OfMRI experiments were performed with 5/10/20 Hz stimulations per each mouse (n = 3 runs per mouse per time-point). The stimulation frequencies were randomized across mice.

